# Downregulation of the secreted protein with an altered thrombospondin repeat (SPATR) impacts the infectivity of malaria sporozoites

**DOI:** 10.1101/2022.03.06.483110

**Authors:** David Mendes Costa, Mónica Sá, Ana Rafaela Teixeira, Begoña Pérez-Cabezas, Sylvain Golba, Hélèna Sefiane-Djemaoune, Pauline Formaglio, Blandine Franke-Fayard, Chris J. Janse, Rogerio Amino, Joana Tavares

**Affiliations:** i3S – Instituto de Investigação e Inovação em Saúde, Universidade do Porto, Porto, 4200-135, Portugal; IBMC – Instituto de Biologia Molecular e Celular, Universidade do Porto, Porto, 4200-135, Portugal; Center for Production and Infection of Anopheles, Institut Pasteur, Paris, 75015, France; Unit of Malaria Infection and Immunity, Institut Pasteur, Paris, 75015, France; Leiden Malaria Research Group, Leiden University Medical Center, Leiden, 2333 ZA, The Netherlands

## Abstract

The identification of surface proteins of the sporozoite stage of malaria parasites important for sporozoite infectivity could aid in the improvement of the efficacy of vaccines targeting pre-erythrocytic stages. Thus, we set out to disclose the role of the secreted protein with an altered thrombospondin repeat (SPATR), which is highly expressed in sporozoites. Previous studies showed an essential function in blood stages, while no role was detected in sporozoites despite high expression. To achieve downregulation of expression in sporozoites while maintaining blood stage expression, a promoter swap approach was used to generate a mutant where the *Plasmodium berghei spatr* gene was placed under transcriptional control of the *hado* gene promoter. Downregulation of expression in oocysts and sporozoites resulted in formation of sporozoites with impaired motility, strongly reduced capacity to invade salivary glands, and decreased infectivity to mice. In conclusion, we revealed a new role for SPATR in sporozoite infectivity, highlighting the importance to use complementary methods in studies on sporozoite biology.

## Introduction

*Plasmodium* sporozoites are motile and invasive stages of the malaria parasites formed inside oocysts in the midgut of the mosquito vector. Fully developed sporozoites egress from the oocysts and are carried by the mosquito circulatory system, the hemolymph, to the salivary glands, which they invade to ensure transmission (1, 2). Once deposited into the skin of the mammalian host by mosquito bite, sporozoites actively migrate and invade blood vessels (3). Sporozoites passively transported by the bloodstream then specifically home to the liver and cross the sinusoidal barrier to get access to hepatocytes, where they multiply (4–6). Thus, sporozoites and the ensuing liver stages mediate the pre-erythrocytic phase of infection, and constitute a transmission bottleneck, as well as ideal vaccination targets (7). The most advanced malaria vaccine, RTS,S/AS01, targets the circumsporozoite protein (CSP), the major sporozoite surface antigen. Anti-CSP antibody titters were shown to be surrogates of protection, but the vaccine had an efficacy of merely 50% or 36% in its target age group against the clinical episodes caused by *Plasmodium falciparum*, the most dangerous malaria parasite to humans, over 1 or 4 years, respectively (8–10).

The development of multivalent subunit vaccines is one of several approaches to improve vaccine efficacy (11). Accomplishing this goal, however, will require greater knowledge of the biology of the parasite, including the identification of sporozoite surface proteins for targeted immune interventions that interfere with the timely completion of the journey of the parasite in the mammalian host. To this end, significant advances have been made to define the *Plasmodium* proteome, including the surface proteome of infectious sporozoites (12–15).

The secreted protein with an altered thrombospondin repeat (SPATR) contains two adhesive domains: an atypical thrombospondin type-1 repeat (TSR) and a Type II epidermal growth factor (EGF)-like domain (16–18). It is conserved across *Plasmodium* species and has several orthologues among the Apicomplexa phylum (19). Although its gene is transcribed in all invasive stages (sporozoites, merozoites and ookinetes) (20, 21), SPATR levels are upregulated in mature sporozoites of both human and rodent infecting *Plasmodium* species (12, 15). Moreover, immunolocalization studies detected the protein at the surface of sporozoites of several *Plasmodium* species (16,18,19), despite the lack of predicted transmembrane domains or a glycosylphosphatidylinositol anchor. Antibodies against the SPATR of *P. falciparum* also blocked hepatocyte invasion by sporozoites (18).

The function of SPATR has been previously addressed using standard reverse genetic approaches. The multiple failed attempts to generate a *spatr* knockout (KO) mutant in *Plasmodium berghei* blood stage parasites, the stage where genetic modification of *Plasmodium* parasites is performed, indicate an essential role for SPATR in blood stages. (19, 22). The stage-specific conditional KO of *spatr* using a flippase-flippase recombination target (Flp-FRT) system (23), which has been used to study gene function in sporozoites (24), consolidated the indispensability of SPATR for the *P. berghei* blood stage (19). However, no defects in sporozoite infectivity to the mosquito or hepatocytes were detected using this conditional KO line (19). In fact, upon hepatocyte infection with sporozoites, merozoites were formed but failed to infect red blood cells (19). In the Flp-FRT system used in this study (19), the excision of the desired genetic sequence, in this case the *spatr* gene, through the stage-specific expression of the Flp recombinase is frequently incomplete at the population level, which may affect to correctly define the phenotype of mutant parasites lacking complete knockout of the gene (19,23,24). Additionally, translational repression of gene expression that has been shown to occur for transcripts of several genes during sporozoite maturation adds another limitation to this Flp-FRT system (15). In fact, gene excision achieved using the thermolabile variant of Flp (FlpL) under transcriptional control of the thrombospondin-related anonymous protein (TRAP) gene promoter mostly occurs in sporozoites after invasion of the salivary glands. This hampers the analysis of genes whose transcripts are already produced during sporozoite formation in the oocysts. Due to these limitations, distinct strategies might be required to ascertain the involvement of SPATR in sporozoite formation and infectivity.

In this study, we confirmed the essentiality of *P. berghei* SPATR for the asexual blood stages and we employed a promoter swap approach to specifically downregulate *spatr* expression early during oocyst development and sporozoite maturation in an attempt to disclose its role in sporozoite infectivity.

## Results

### Knockdown of *spatr* in *P. berghei* sporozoites

We attempted to knock out the *spatr* gene by double crossover homologous recombination, using two different strategies (Fig. S1). We first made use of a *Plasmo*GEM (Sanger) targeting vector (Fig. S1A), which contains long homology regions that increase the efficiency of integration of the construct at the target locus (29, 38). While integration at the correct site was confirmed in transfectant populations, isogenic transgenic parasites could not be isolated in several independent attempts (Fig. S1B). To exclude the possibility that the long homology regions that span over neighboring genes could have also led to alterations in the expression of these genes, affecting the viability of mutant parasites, we constructed a different transfection vector that restricted the homology regions to the intergenic sequences flanking the *spatr* gene (Fig. S1C). However, we were unable to detect the presence of transgenic parasites within the parental populations that arose after the transfection by PCR (data not shown). These results support other reports of SPATR essentiality for *P. berghei* replicative blood stages (19, 22).

Thus, to ascertain the potential involvement of SPATR in sporozoite infectivity to the mosquito and mammalian hosts, we devised a promoter swap strategy to attain the knockdown of SPATR expression early during sporozoite development. In this strategy, the *spatr* gene is placed under transcriptional control of a promoter of a gene that is active in blood stages but is turned off in sporozoites. Available RNA-seq datasets showed higher abundance of *spatr* transcripts in blood schizonts, ookinetes and salivary gland sporozoites (20, 21) (Fig. 1A). Based on these criteria, we searched genome-wide RNA-seq data and selected the genes that encode the haloacid dehydrogenase domain ookinete protein (HADO) and the putative centrosomal protein 76 (CEP76; Fig. 1A and S2A). Both genes are transcribed in blood stages and in ookinetes. While the absence of *hado* transcripts in oocysts had been previously reported (39), no such data was available for the *cep76* gene, although we hypothesized this gene would be expressed in oocysts during sporozoite formation due to its expected role in cell division (40). We selected this gene to use its promoter a control to investigate the possible role for translational repression of transcripts in sporozoites (15; and see Introduction section). To confirm the RNA-seq expression data, we evaluated the transcriptional profiles of *spatr* and the selected genes by RT-qPCR (Fig. 1B and S2B). These data show that unlike *spatr*, whose transcription upregulation within oocysts appeared to roughly coincide with the start of sporozoite formation 8 days after the infectious blood meal (41) and peaked in hemolymph sporozoites, *hado* was lowly expressed throughout oocyst development and sporozoite maturation (Fig. 1B). On the other hand, *cep76* transcripts were detected in oocysts but were drastically reduced in hemolymph and salivary gland sporozoites (Fig. S2B).

**Fig 1.**
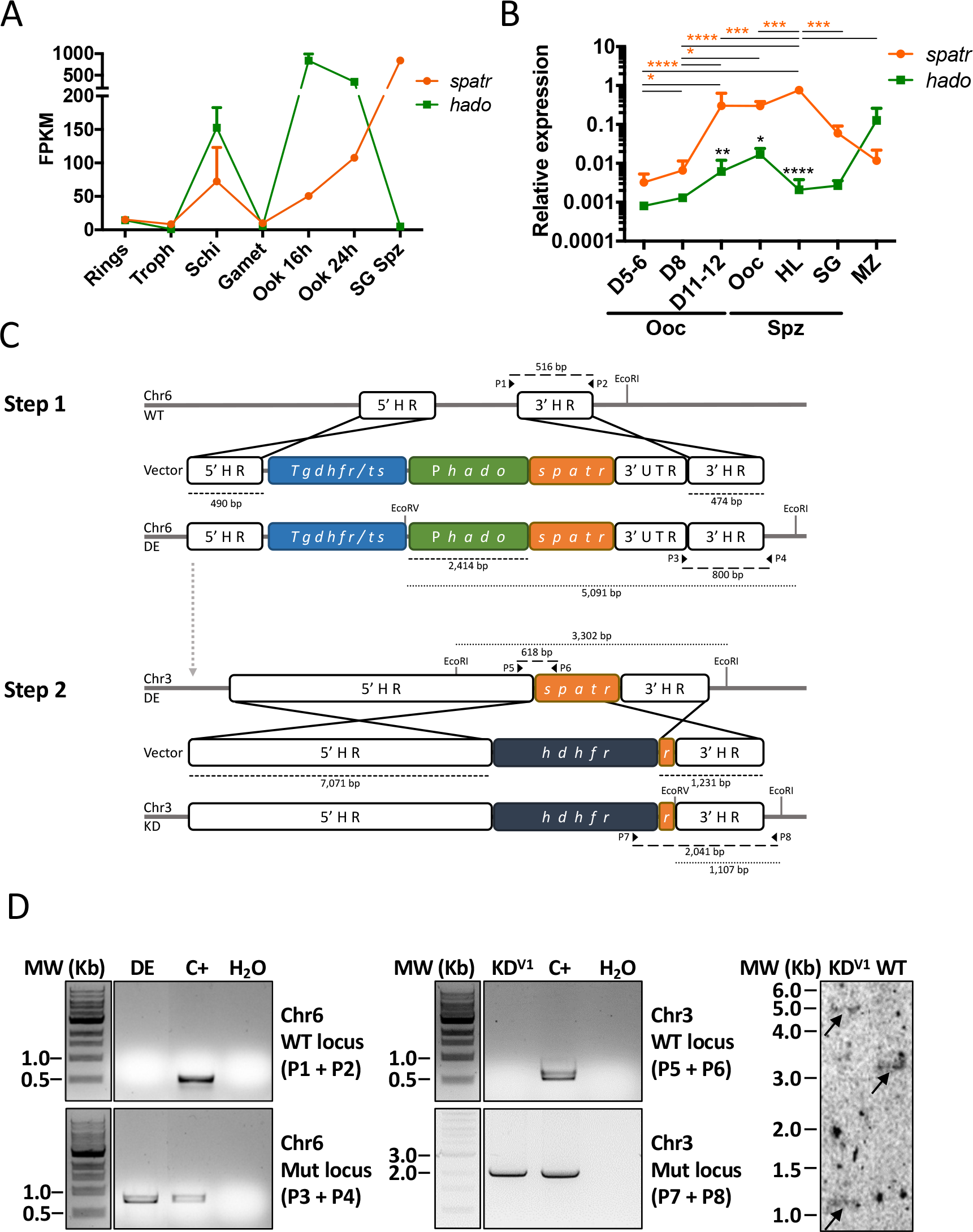
Generation of SPATR knockdown (KD) lines. (A) RNA-seq data of *spatr* and *hado* genes in different stages of the *P. berghei* life cycle. Plotted data was gathered from published datasets [REFS]. (B) Expression of *spatr* and *hado* relative to *hsp70* in different developmental stages and days post-infection, assessed by RT-qPCR (mean values + standard deviation of at least two biological replicates). Statistical analyses of differences between genes (black asterisks) or developmental stages (colored asterisks) were performed using two-way ANOVA with Sidak’s multiple comparison test or two-way ANOVA with Turkey’s multiple comparison test, respectively. Non-significance is not represented. (C) Stepwise schematic representation of the two-step promoter exchange strategy using the promoter of *hado*. The sizes of genomic regions are shown next to dashed line. Sequences are not to scale. (D) Genotyping of transfected populations by PCR (left and center) and Southern blot (right). Restriction sites and primers used for the genotypic analyses by Southern blot or PCR, respectively, are depicted in panel C. Amplicon sizes and sequences detected by Southern blot are shown next to long dashes or dotted lines, respectively. bp: base pairs. C+: positive control (WT or transfer population on the top or bottom gels, respectively). Chr: chromosome. DE: SPATR double-expressor. Gamet: gametocytes. FPKM: fragments per kilobase of transcript per million mapped reads. *hdhfr*: human dihydrofolate reductase. HL: hemolymph. HR: homology region. Kb: kilobases. KD^V1^: SPATR knockdown V1 clone. Mut: mutant. MW: molecular weight. Ooc: oocyst(s). Ook: ookinetes. Schi: schizonts. SG: salivary gland. Spz: sporozoites. *Tgdhfr/ts*: *Toxoplasma gondii* dihydrofolate reductase-thymidylate synthase. Troph: trophozoites. UTR: untranslated region.

The exact location of the promoter elements required to drive the expression of *spatr* in the different life cycle stages is unknown. Therefore, the possibility exists that swapping the short intergenic sequence that precedes the *spatr* gene by the two selected promoters affects 3’ regulatory elements of the upstream gene. Moreover, neighboring *Plasmodium* genes can form co-expressed units, which may be controlled by a single promoter (42), and bicistronic transcripts have been reported in *P. berghei* (43). *Plasmodium* also utilizes several layers of tight transcriptional control, some of which are promoter-independent (44). Thus, changing the promoter of the *spatr* gene in its native locus could still lead to transcription in the mosquito stages regardless of the stage-specificity of the new promoters. For this reason, we opted for a conservative promoter exchange approach that involved the insertion of a second copy of *spatr* under transcriptional control of the *hado* or *cep76* putative promoters in a previously reported silent locus in chromosome 6 (30), and the subsequent deletion of the endogenous *spatr* gene located in chromosome 3 (Fig. 1C and S2C). The generated intermediate lines were named SPATR double-expressers (SPATR DE), as they were expected to possess two active copies of the gene in several developmental stages. While we could delete the endogenous *spatr* gene in the SPATR DE with the *hado* promoter (SPATR KD parasites; 2414 bp; Fig. 1C–D), we failed to delete the endogenous *spatr* gene in the SPATR DE with the *cep76* promoter (845 bp; Fig. S2D). This may be due to insufficient or ill-timed expression of *spatr* in blood stages under control of the *cep76* promoter. Our RT-qPCR analysis indicates that *spatr* transcripts in blood stages (merozoites) of WT parasites are approximately two times more abundant than *cep76* transcripts, although the difference was not statistically significant (Fig. S2B). Alternatively, we cannot exclude that the selected *cep76* 5’ intergenic region does not contain the promoter elements that drive the expression of the gene. Therefore, references to SPATR DE and SPATR KD parasites will henceforth only refer to the isogenic lines containing a *spatr* copy under control of the *hado* promoter.

### SPATR is important for the colonization of the mosquito salivary glands

SPATR KD blood stages displayed normal growth (Fig. 2A) and successfully differentiated into functional gametocytes (Fig. 2B), as evidenced by the formation of microgametes (Fig. 2C) and their ability to infect mosquitos (Fig. 2D). While the SPATR DE line showed a prevalence of infected mosquitos (Fig. 2D) and sporozoite numbers (Fig. 2E) comparable to the WT, salivary glands of SPATR KD-infected mosquitoes showed strongly reduced sporozoite numbers (Fig. 2E–F). Interestingly, the reduced numbers of sporozoites in the salivary glands did not result in the accumulation of sporozoites in the hemolymph (Fig. 2E). Few SPATR KD sporozoites were detected within the salivary glands and these did not show obvious structural modifications by transmission electron microscopy analysis (Fig. 2F). This indicates that some parasites were indeed capable of infecting the mosquito salivary glands, rather than remaining associated to the basal lining of the glands. Although we only observed a very small number of SPATR KD sporozoites inside the salivary glands, no parasites were seen within the secretory cavities. Conversely, high numbers of WT sporozoites were present both in the parenchyma and in the secretory cavities of the salivary glands (Fig. 2F).

**Fig 2.**
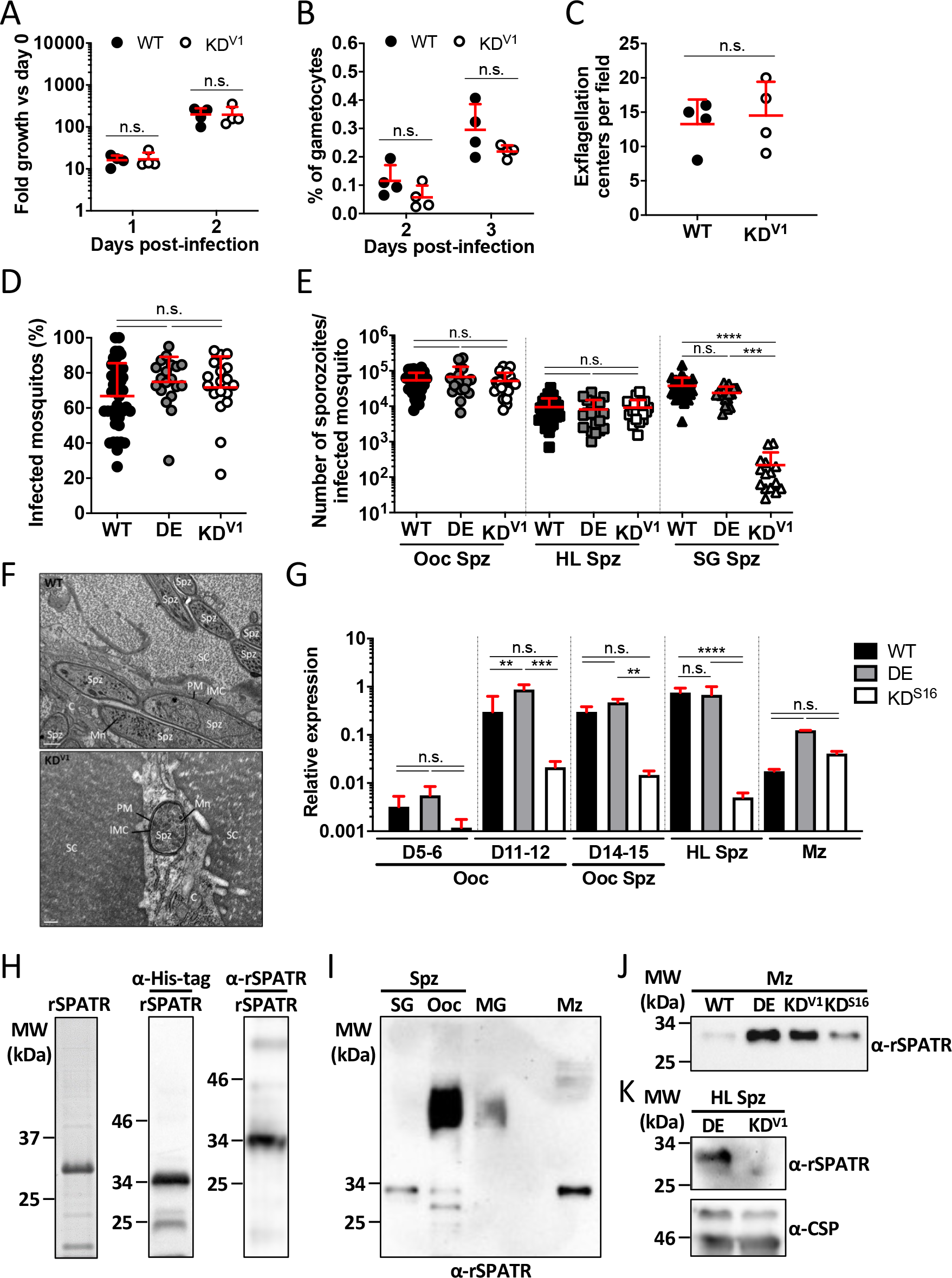
Infectivity of SPATR KD parasites in the blood and mosquito, and evaluation of *spatr* expression. (A) Fold-growth of blood stages versus parasitemia on infection day. (B) Gametocytemia. (C) Microgamete exflagellation. (D) Prevalence of infected mosquitos in independent experiments. (E) Numbers of oocyst, hemolymph and salivary gland sporozoites per mosquito in independent experiments. (F) Ultrastructure and localization of sporozoites within the salivary glands. Scale bars: 500 nm (top) or 200 nm (bottom). (G) Expression of *spatr* relative to *hsp70* in WT, SPATR DE and SPATR KD parasites determined by RT-qPCR using at least two biological replicates. (H) SDS-PAGE gel showing 5 µg of purified recombinant SPATR (rSPATR; left), and reactivity of an α-His-tag antibody (center) and the mouse α-SPATR polyclonal antibodies (right) against 100 ng of rSPATR. (I) Detection of SPATR by Western blot in samples prepared with 1.25×10^4^ WT salivary gland and oocyst sporozoites or 1.5×10^6^ WT merozoites. An amount of uninfected midguts (MG) equivalent to that loaded in the oocyst sporozoite lane was used to test the reactivity of the polyclonal antibodies against mosquito-derived material. (J) Western blot analysis of merozoite extracts prepared with 4×10^6^ merozoites of the different lines. (K) SPATR levels in 1.5×10^4^ SPATR DE or SPATR KD hemolymph sporozoites determined by Western blot. CSP levels evaluated with mouse anti-CSP monoclonal antibody 3D11 were used as a loading control. Mean values + standard deviation are shown in panels A–E and G. Statistical analyses of were performed using the Mann-Whitney test (A–C) or the Kruskal-Wallis test with Dunn’s multiple comparisons test (D, E, G). n.s.: non-significant. C: salivary gland cell. D: day post-infection. DE: SPATR double-expressor. HL: hemolymph. IMC: inner membrane complex. KD^S16^: SPATR knockdown S16 clone. KD^V1^: SPATR knockdown V1 clone. MW: molecular weight. Mn: microneme. Mz: merozoites. Ooc: oocyst(s). PM: plasma membrane. SC: secretory cavity. SG: salivary gland. Spz: sporozoites.

To confirm the decreased transcription of *spatr* in SPATR KD sporozoites, we analyzed oocysts at different time points after the infectious blood meal, sporozoites collected from mechanically disrupted oocysts, hemolymph sporozoites and purified merozoites by RT-qPCR (Fig. 2G). Transcripts were found to be significantly less abundant in SPATR KD oocysts and sporozoites from days 11–12 post-infection than in the control lines. On the other hand, the SPATR DE line had similar levels of *spatr* transcripts in comparison to the WT line in all sporozoite stages, although transcript levels were around 7 times higher in SPATR DE merozoites (Fig. 2G). This is indicative that, as expected, both copies of the *spatr* gene are being transcribed in the merozoite stage (Fig. 2G). Mouse polyclonal antibodies against a His-tagged recombinant SPATR (rSPATR; Fig. 2H) detected bands of the approximate predicted size of SPATR (29 or 26 kDa with or without the signal peptide, respectively) in extracts of sporozoites collected from the salivary glands and oocysts, as well as in merozoites (Fig. 2I). As anticipated, SPATR was also expressed in SPATR DE and SPATR KD merozoites (Fig. 2J), but protein levels were drastically reduced in SPATR KD hemolymph sporozoites (Fig. 2K). The detection of the protein in WT oocyst-derived sporozoites was unexpected, since previous proteomics studies performed in *P. falciparum* and *P. yoelii* had only detected SPATR protein in salivary gland sporozoites, whereas transcripts were present in sporozoites collected from both the oocysts and the salivary glands, which was suggestive of translational repression^12,15^. Whether this was due to differences in the parasite species, the technical approaches, or the age of oocyst sporozoites – these were collected from day 19 post-infection onwards in this work, as opposed to days 10–14 in the proteomics studies – remains to be elucidated. Similar band patterns were obtained by Western blot using extracts of sporozoites expressing a c-Myc-tagged SPATR at the C-terminus (Fig. S3). Moreover, the tagging of the protein was not detrimental to the parasite as SPATR-myc sporozoites were present in similar numbers compared to controls in oocysts and in the salivary glands (Fig. S3D) and no defects were observed during blood stage growth in any of the infections resulting from the generation and testing of the isogenic lines (data not shown).

### Kinetics of SPATR KD sporozoite release to the hemolymph are altered

In an attempt to understand why the reduced numbers of salivary gland sporozoites did not result in the accumulation of SPATR KD sporozoites in the hemolymph, we evaluated oocyst and sporozoite development from early time points post-infection (Fig. 3A–E). Neither the number (Fig. 3A) nor the size (Fig. 3B) of the oocysts revealed any clear developmental impairments. However, whereas the number of sporozoites per oocyst did not vary significantly in the WT and SPATR DE lines from day 11 until day 15 post-infection, a significant increase in the number of sporozoites per oocyst was observed in both SPATR KD isogenic lines on day 15 (Fig. 3C). When the same ratio was applied to hemolymph sporozoites, all lines exhibited an overall similar increase in numbers over time (Fig. 3D). The lower numbers of sporozoites inside oocysts on days 11–13 post-infection could be met with a transient accumulation of the few parasites released from oocysts in the hemolymph, resulting in similar sporozoite numbers in the circulatory system of the mosquito. This hypothesis is supported by the observation that the mortality of SPATR KD hemolymph sporozoites was not superior to that of WT parasites (Fig. 3F) and that sporozoites were not being sequestered in other tissues of the mosquito (Fig. 3G). Indeed, SPATR KD oocysts also contained more sporozoites than the control lines from day 24 post-infectious blood meal (Fig. 3H). Altogether, these data could indicate a defect in egress of sporozoites from the oocyst.

**Fig 3.**
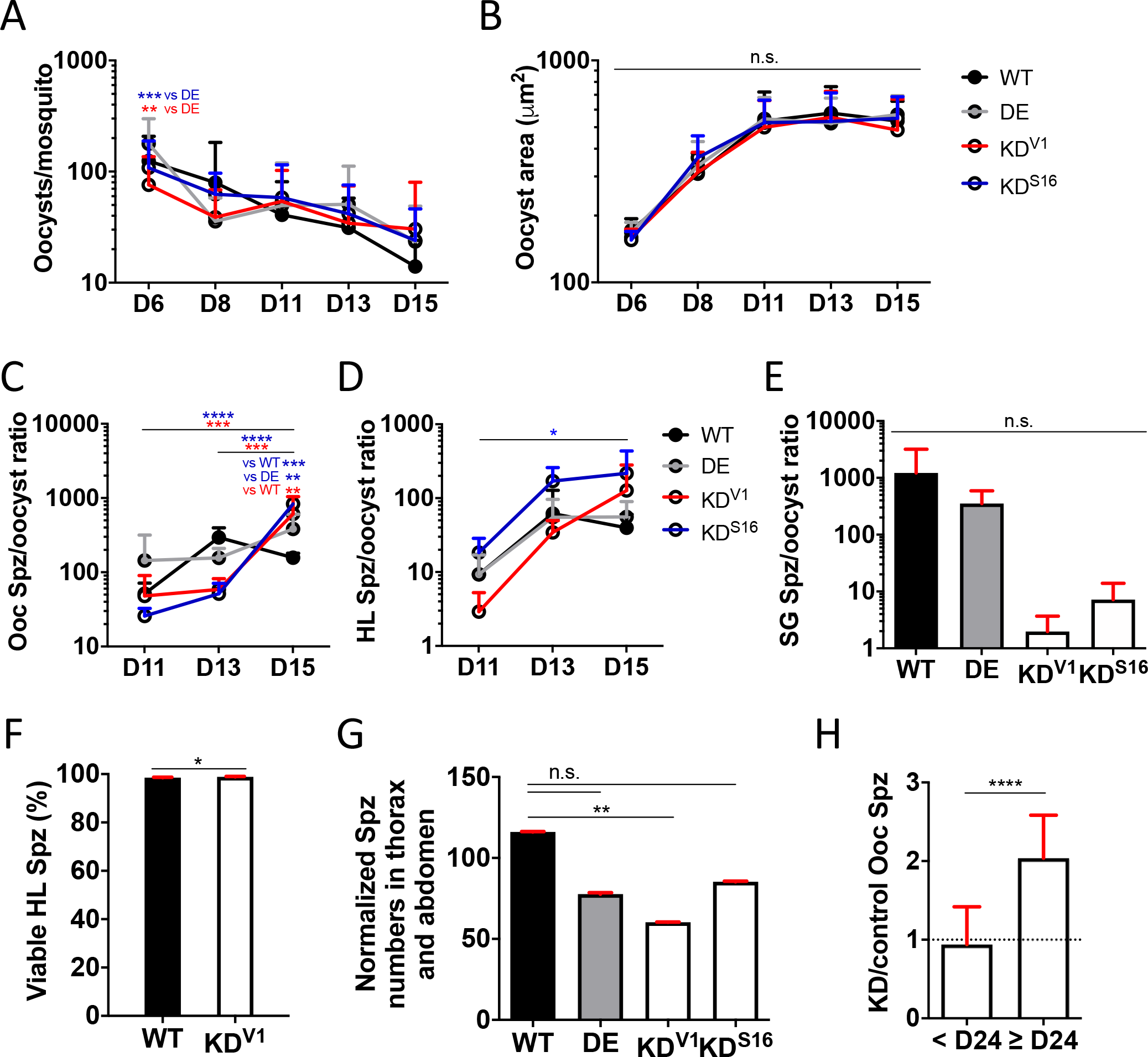
Analysis of the development and location of SPATR KD parasites in the mosquito. (A) Schematic representation of the generation of the SPATR-myc line, which expressed SPATR fused to two c-myc sequences (myc) at C-terminal. The duplicated copy of the gene at 3’ contains a mutation in the start codon (ACG) to prevent the expression of the full length and untagged SPATR. The size of the homology region is shown below the dashed line. Sequences are not to scale. (B) Genetic analysis of two clonal populations by PCR. Genomic DNA from WT parasites or a transfer population (TP) were used as positive controls (top and bottom gels, respectively), while SPATR KDV1 was used as a positive and controls. The primers used for the genetic analysis and the amplicon sizes (long dashes) are depicted in panel A. (C) Reactivity of α-c-Myc and α-rSPATR antibodies to extracts prepared with 4×105 SPATR-myc oocyst sporozoites or 1.5×105 SPATR-myc salivary gland sporozoites, evaluated by western blot. (D) Numbers of SPATR-myc (Myc) oocyst, hemolymph and salivary gland sporozoites per mosquito. WT values are also shown in Fig. 2E. Mean values + standard deviation of independent determinations are represented. Statistical analysis was performed using the Mann-Whitney test (F and H) or the Kruskal-Wallis test with Dunn’s multiple comparisons test (A–E and G). n.s.: non-significant (not shown on panels A, C and D). bp: base pairs. Chr3: chromosome 3. HL: hemolymph sporozoites. IR: intergenic region. Mut: mutant. MW: molecular weight. SPATR My Ooc: oocyst. SG: salivary gland. Spz: sporozoites. *Tgdhfr/ts*: *Toxoplasma gondii* dihydrofolate reductase-thymidylate synthase.

We also performed a qualitative analysis of CSP and TRAP expression in SPATR KD hemolymph sporozoites, as both these proteins play a role in invasion of the salivary glands (45–58). No overt differences in expression and location of CSP and TRAP were detected (Fig. S4). The surface staining of CSP also allows a crude inspection of the morphology of the sporozoite, which was not overtly different from that of WT parasites (Fig. S4A). The typical location of TRAP showed that the biogenesis, subcellular location and discharge of TRAP-containing micronemes remained unaffected (Fig. S4B).

### The establishment of liver infection in mice by SPATR KD sporozoites is impaired

To evaluate sporozoite infectivity, parasites were collected from the hemolymph and injected intravenously in mice. While SPATR DE parasites successfully accomplished the liver stage and subsequent blood infections, parasite liver loads in SPATR KD-infected mice were strongly reduced (as shown by close to background level bioluminescent signals in the liver) and were rarely able to establish a blood infection (Fig. 4A and Table S2). A milder defective phenotype was observed when we infected mice with the few SPATR KD sporozoites that we were able to collect from salivary glands, with reduced parasite liver loads and prolonged prepatent periods of approximately 1 day (Fig. 4B and Table S2). Conversely, infections with SPATR DE sporozoites consistently yielded similar parasite liver loads and prepatency to those obtained with the WT line (Fig. 4A–B and Table S2).

**Fig 4.**
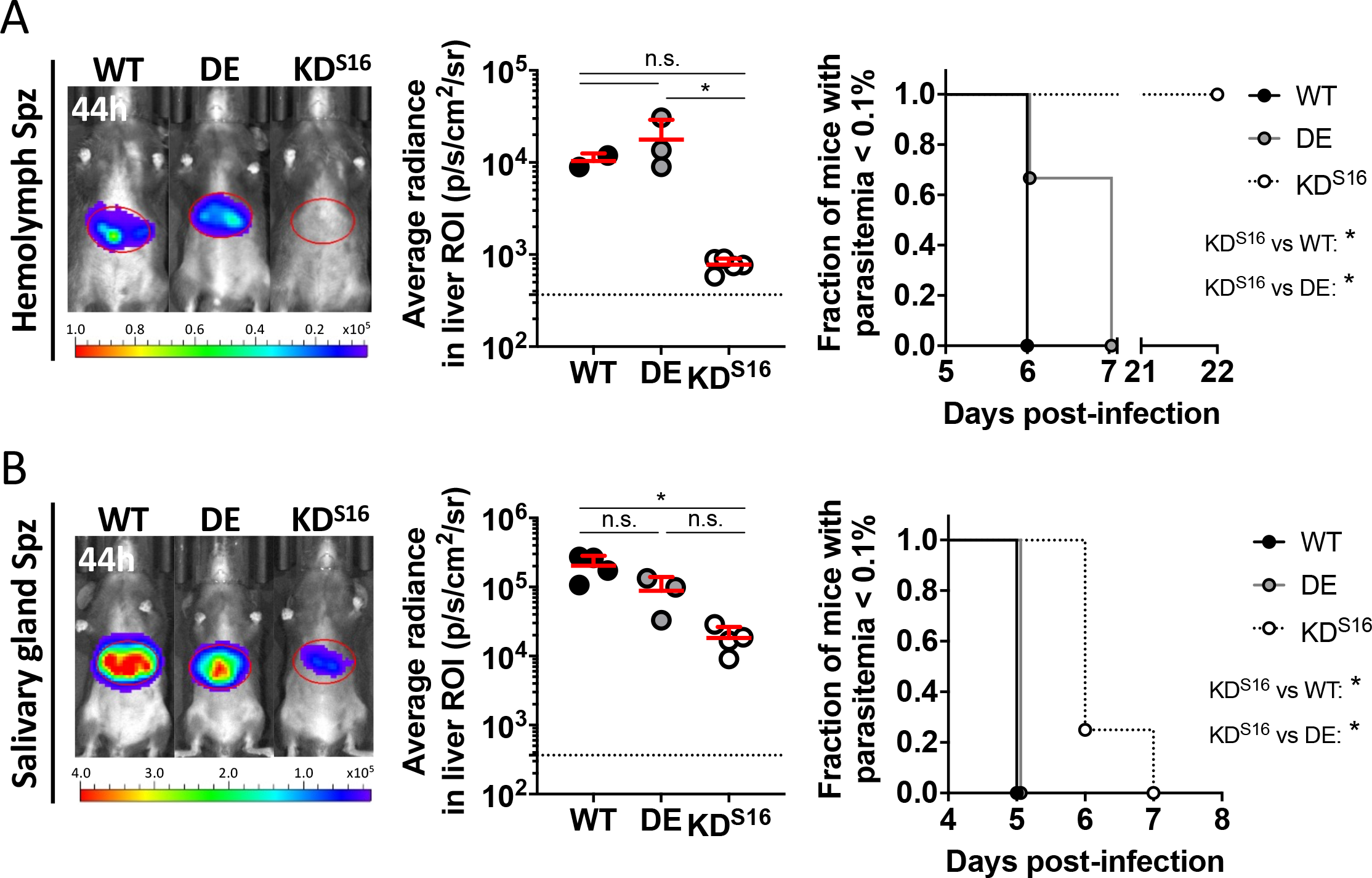
Infectivity of SPATR KD sporozoites. (A, B) Infection of C57BL/6 mice with 2×10^4^ hemolymph (A) or 2,500 salivary gland sporozoites (B). A representative image resulting superimposition of the bioluminescence signal map and a grey-scale photograph of the mice at 44 h post-infection (left), the quantification of the signal in the defined liver ROI (center), and the fraction of mice with parasitemia above 0.1% (right) are shown. Dotted lines represent the average radiance of uninfected mice. Representative experiments are shown. The mean values + standard deviations are shown in both center panels. Statistical analyses were performed using the Kruskal-Wallis test with Dunn’s multiple comparisons test (center) or the Gehan-Breslow-Wilcoxon test corrected for multiple comparisons (right). n.s.: non-significant. DE: SPATR double-expressor. KD^S16^: SPATR knockdown S16 clone. Spz: sporozoite.

### SPATR KD sporozoites exhibit defects in hepatocyte invasion, migration and homing to the liver

A decrease in parasite liver loads two days after intravenous injection of SPATR KD sporozoites can be attributable to impairments in parasite arrest in the liver sinusoids, crossing of the sinusoidal barrier, recognition and invasion of hepatocytes, or development within host cells (5, 7). Thus, we first evaluated the capacity of SPATR KD parasites to invade and infect hepatocytes *in vitro* using the HepG2 hepatoma cell line. DE hemolymph sporozoites did not display any infectivity or developmental defects *in vitro* (Fig. 5A–C). Conversely, invasion assays performed with SPATR KD hemolymph sporozoites yielded at least 5 times fewer intracellular sporozoites than with the WT line (Fig. 5A), which also resulted in lower numbers of SPATR KD exoerythrocytic forms (EEFs) 24 to 48 h post-infection (Fig. 5B). While variations in EEF size (205 ± 79 µm^2^ vs 166 ± 79 µm^2^, for WT and SPATR KD, respectively) were observed (Fig. 5C), which could indicate a developmental impairment, the small numbers of SPATR KD parasites analyzed preclude the definite assignment of a defect in liver stage development to SPATR KD parasites.

**Fig 5.**
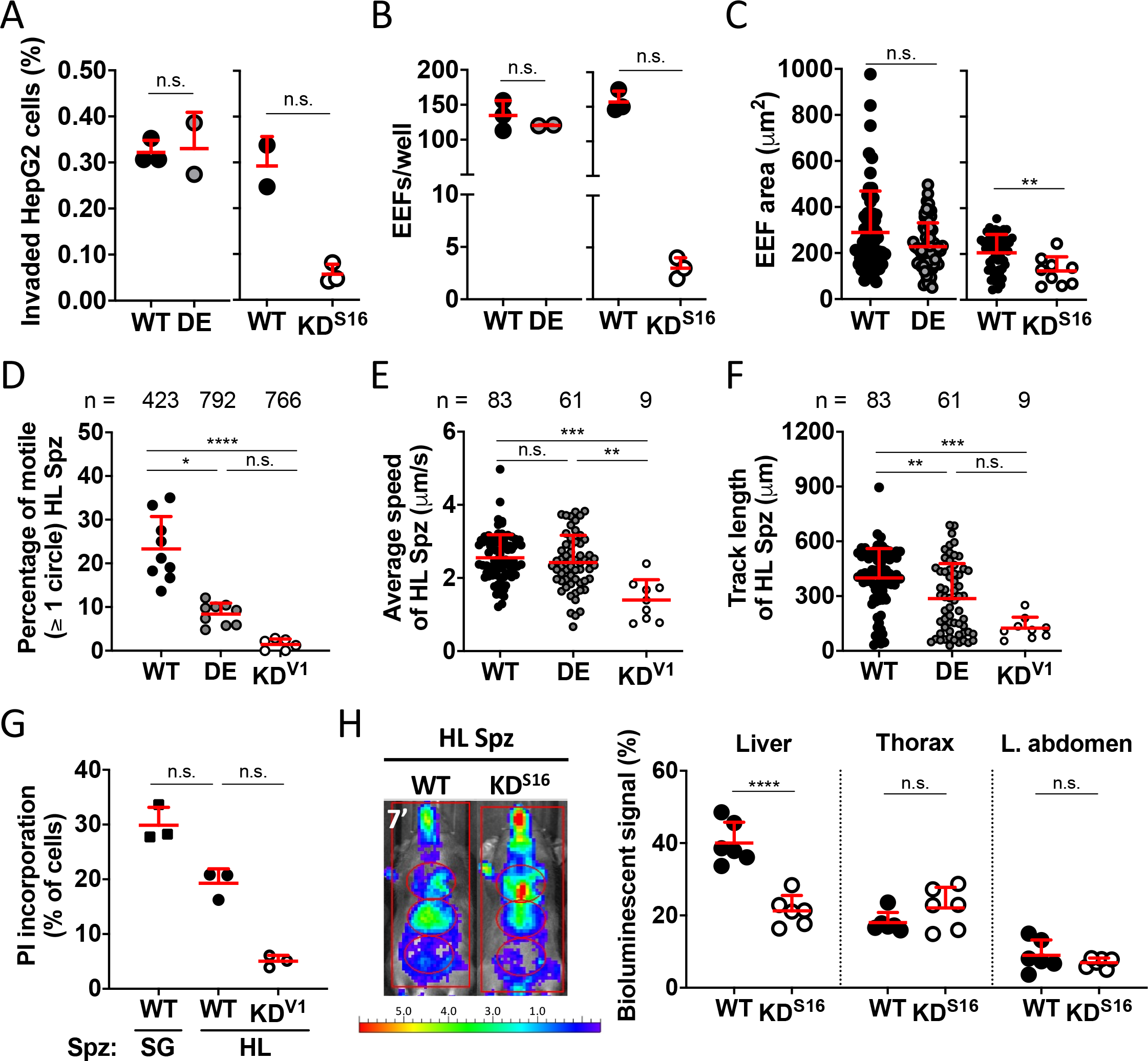
*In vitro* and *in vivo* studies with SPATR KD hemolymph sporozoites. (A) Invasion of HepG2 cells by hemolymph sporozoites. (B, C) Numbers (B) and area (C) of EEFs following the infection of HepG2 cells with hemolymph sporozoites (D–F) Percentage of motile hemolymph sporozoites (D), and their average speed (E) and track length (F). Numbers above the plots indicate the total number of sporozoites analyzed. The symbols in panel D represent the percentages obtained in different videos, rather than individual sporozoites analyzed. (G) Determination of PI incorporation by HepG2 cells by flow cytometry analysis following incubation with sporozoites. The average percentage of PI+ cells only incubated with PI (no sporozoites) was subtracted to the percentage of PI+ cells obtained in each experimental condition. WT salivary gland sporozoites were used as a positive control. (H) Homing of hemolymph sporozoites from 7 minutes post-infection. A representative superimposed image is shown on the (left) next to the percentage of total photons detected in the several anatomical regions of C57BL/6 mice (right). Data from two independent experiments were pooled. Mean values + standard deviations are represented and representative experiments are shown (except in panel H). Statistical analyses were performed using the Mann-Whitney test (A–C, G–H) or the Kruskal-Wallis test with Dunn’s multiple comparisons test (D–F). n.s.: non-significant. All panels show data from representative experiments. DE: SPATR double-expressor. HL: hemolymph. KD^S16^: SPATR knockdown S16 clone. KD^V1^: SPATR knockdown V1 clone. SG: salivary gland. Spz: sporozoites.

Gliding motility and cell traversal activity allow *Plasmodium* sporozoites to travel through tissues and reach their destination in the liver. Therefore, we then investigated the motility of SPATR KD sporozoites in a 2D system. We observed a marked reduction in the percentage of gliding SPATR KD hemolymph sporozoites (Fig. 5D), together with a decrease in speed (Fig. 5E), which was accompanied by an inferior total travel distance in comparison to the WT line (Fig. 5F). The decrease in the proportion of motile SPATR KD sporozoites was accompanied by an increase in the percentage of parasites bound to the substrate but not exhibiting productive motility (Fig. S5A) and the total percentage of adhered parasites remained similar in all lines (Fig. S5B). Thus, SPATR KD hemolymph sporozoites appear to retain the ability to interact with the coverslip. These data suggest the strongly reduced *in vivo* infectivity of SPATR KD sporozoites could, in part, stem from their impaired motility. Interestingly, we also observed fewer SPATR DE hemolymph sporozoites gliding (Fig. 5D) and motility was sustained for shorter periods of time than for the WT line, as demonstrated by the reduced track lengths despite normal speed (Fig. 5E–F). However, these differences did not significantly impact the infectivity of SPATR DE hemolymph sporozoites *in vivo* and *in vitro* (Fig. 4A and 5A–C) and were not observed when we compared the motility of DE and WT salivary gland sporozoites (Fig. S5C–E).

Next, we tested the ability of SPATR KD hemolymph sporozoites to wound and traverse host cells, which was measured by the incorporation of propidium iodide, a membrane-impermeable intercalating agent, by hepatocytes following incubation with the parasites. The modest reduction in cell wounding activity of SPATR KD hemolymph sporozoites (1.4- to 3.6-fold, in two independent experiments; Fig. 5G) indicates that cell traversal should play a minor role in the severely impaired infectivity of this mutant line in mice.

Finally, to dissect the capacity of SPATR KD sporozoites to home to the liver, we infected mice with hemolymph sporozoites and quantified the bioluminescent signal shortly after intravascular injection. Importantly, it has been shown that Kupffer cells function as facultative entry points to the liver parenchyma for salivary gland sporozoites (4, 59). However, hemolymph sporozoites do not possess a fully-matured cell traversal machinery (60) and may be retained by Kupffer cells, as was observed for the cell traversal-deficient sporozoites lacking the sporozoite protein essential for cell traversal (SPECT2) (4). Therefore, homing experiments were performed in clodronate-treated mice for Kupffer cell depletion to ensure the detected signal more accurately reflected parasite capacity to be arrested in the liver sinusoids. In contrast to the partial accumulation of WT hemolymph sporozoites in the anatomical area corresponding to the liver (Fig. 5H), the percentage of bioluminescent signal in the liver of mice infected with SPATR KD hemolymph sporozoites (21%; Fig. 5H) was similar to those obtained in mice infected with circulating parasites such as *Trypanosoma brucei* bloodstreams forms or *P. berghei* merozoites (6). These data show that SPATR KD hemolymph sporozoites fail to accumulate in their target organ in this short time frame.

In conclusion, these cumulative impairments explain in the incapacity of these sporozoites to productively infect the liver.

### Genetic complementation partially reverts the SPATR KD phenotype

In an attempt to correlate the loss of infectivity observed in SPATR KD sporozoites to the decrease in SPATR levels, we genetically complemented the SPATR KD V1 isogenic line with an episome carrying part of the *spatr* locus (Fig. 6A). Since the nature and location of the promoter elements required for the transcription of *spatr* have not yet been identified, we also constructed a plasmid containing the *spatr* open reading frame downstream of the characterized promoter of *trap* (Fig. 6A), which has been shown to drive the expression of genes in ookinetes and sporozoites starting from the oocyst stage (23, 61). Moreover, we confirmed that the transcriptional profile of *trap* in selected stages of the parasite was similar to that of *spatr* (Fig. 6B). The plasmid also contained the centromeric region of chromosome 5 of *P. berghei*, the PbCEN5-core, to enhance segregation stability during cell division (31–33). Lastly, the RedStar red fluorescent protein (RFP) reporter gene present in the centromeric plasmid allowed us to perform the enrichment RFP-positive blood stage parasites by FACS (Fig. 6C). While erythrocytes harboring parasites complemented with the episome containing the *spatr* gene under transcriptional control of the promoter of *trap* (SPATR COMP^P*trap*^) represented approximately half of the total infected erythrocyte population after several enrichment steps, the percentage of erythrocytes carrying parasites transfected with the version of the plasmid bearing the putative promoter of *spatr* (SPATR COMP^P*spatr*^) never surpassed 10% (Fig. 6C).

**Fig 6.**
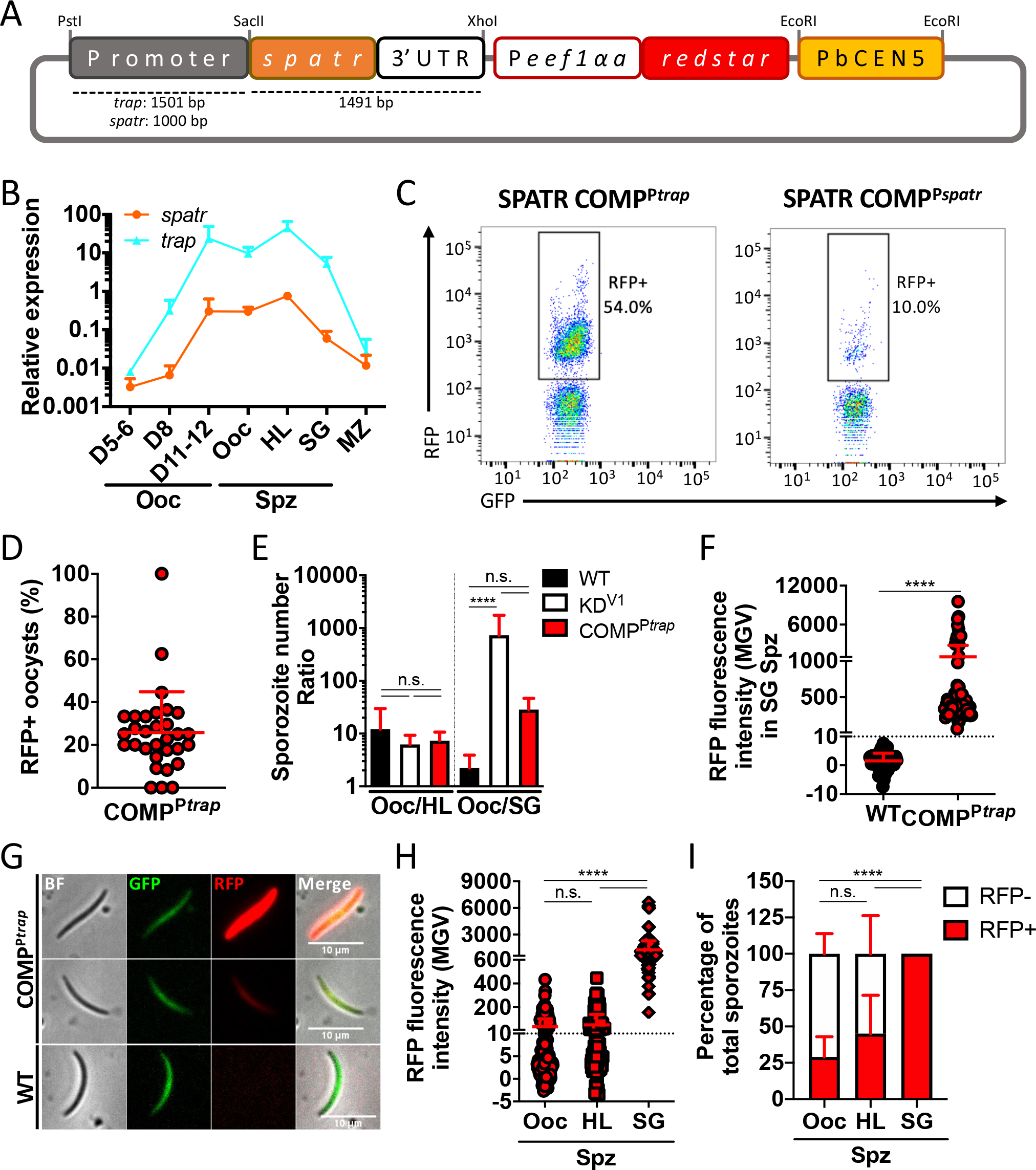
Genetic complementation of SPATR KD parasites. (A) Centromeric episome used to complement the SPATR KD^V1^ line. Restriction enzymes and size of the sequences (not to scale) used to construct the plasmid are indicated. (B) Expression of *trap* relative to *hsp70* in different developmental stages and days post-infection, assessed by RT-qPCR using at least two biological replicates. The transcription data of *spatr* shown in Fig. 1B were plotted here to facilitate comparison of the transcriptional profiles. (C) FACS analysis of blood collected from mice infected with either SPATR COMP^P*trap*^ (left) or SPATR COMP^P*spatr*^ (right) populations. The percentages of parasites harboring the episome (RFP-positive events gated in the GFP-positive population) are shown. (D) Percentage of RFP-expressing oocysts at day 15 post-infection, determined by fluorescence microscopy. Each symbol represents an individual midgut. (E) Ratios of oocyst to either hemolymph or salivary gland sporozoite numbers, from days 14 to 26 post-infection. (F) Fluorescence intensity of live salivary gland sporozoites on day 26 post-infection. The dotted line represents the maximum fluorescence intensity value obtained for WT salivary gland sporozoites with the same image acquisition settings. (G) Representative immunofluorescence images of gland sporozoites on day 26 post-infection. (H) RFP fluorescence intensity of live oocyst, hemolymph and salivary gland SPATR COMP^P*spatr*^ sporozoites on day 19 post-infection. The dotted line represents the maximum fluorescence intensity value obtained for WT salivary gland sporozoites with the same image acquisition settings (F). (I) Percentage of RFP-expressing SPATR COMP^P*spatr*^ sporozoites, using the cutoff established on panels F and H. Representative experiments are shown and mean values + standard deviation are represented. Statistical analyses were performed using the Mann-Whitney test (F) or the Kruskal-Wallis with Dunn’s multiple comparisons test (E, H–I). n.s.: non-significant. BF: brightfield. bp: base pairs. HL: hemolymph. MGV: mean grey values. MZ: merozoite. Ooc: oocyst(s). P*eef1αa*: promoter of the *eef1αa* gene. SG: salivary gland. Spz: sporozoites. UTR: untranslated region.

Therefore, we only subjected the mixed populations containing SPATR COMP^P*trap*^ parasites to a mosquito passage. The average percentage of RFP-positive oocysts (26%; Fig. 6D) was lower than the percentage of RFP-positive blood stages (54%; Fig. 6C), which implies further loss of the episome during schizogony prior to the infectious blood feeding or during multiplication within the oocysts. Nonetheless, the genetic complementation resulted in the increase of sporozoites inside the salivary glands in mosquitos infected with SPATR COMP^P*trap*^ parasites, as demonstrated by the reduced ratio between the numbers of oocyst and salivary gland sporozoites in comparison to the SPATR KD line (Fig. 6E). Despite the fact that only nearly 1 in 4 SPATR COMP^P*trap*^ oocysts contained the plasmid (Fig. 6D), which could partially explain why we did not attain full reversion of the defective salivary gland invasion phenotype (Fig. 6E), almost all SPATR COMP^P*trap*^ sporozoites present in the salivary glands had detectable RFP signal (Fig. 6F–I). Moreover, the proportion of sporozoites carrying the plasmid also increased from the oocyst and hemolymph sporozoite populations to the parasite population present in the salivary glands (Fig. 6I). This demonstrates that parasites carrying the episome specifically accumulate in the salivary glands and links the low numbers of SPATR KD salivary gland sporozoites to the downregulation of *spatr* expression.

We then attempted to determine if the genetic complementation of the SPATR KD parasites restored sporozoite infectivity to mice. The injection of 60,000 SPATR COMP^P*trap*^ hemolymph sporozoites in two mice resulted in the detection of parasites in the blood of both animals, while none of the 5 mice injected with the same inoculum of SPATR KD became infected (Table S2). This is consistent with a recovery of hemolymph sporozoite infectivity, especially if we consider 45% of COMP^P*trap*^ hemolymph parasites carry the episome, meaning the starting inoculum corresponds to ∼27,000 RFP-positive sporozoites (Fig. 6I). Interestingly, the percentage of RFP-positive blood stages in the populations arising from these infections, as well as infection initiated with COMP^P*trap*^ salivary gland sporozoites (Table S2), ranged from 0-10%. Since SPATR KD hemolymph sporozoites rarely managed to infect mice and close to 100% of SPATR COMP^P*trap*^ salivary gland sporozoites were determined to express RFP, it is likely that these infections were initiated with RFP-expressing sporozoites, but the plasmid was lost during schizogony, supporting out previous conclusions.

## Discussion

SPATR is a conserved protein with adhesive domains expressed in all invasive stages of *Plasmodium*. Despite the upregulation of transcription in infectious sporozoites, we found that SPATR is, as previously described, essential for the asexual blood stages (12,15,19). Using a promoter swap approach to specifically downregulate *spatr* expression early during sporozoite development, we established that SPATR is required for the invasion of the mosquito salivary glands by sporozoites. Moreover, the ability of SPATR KD sporozoites collected from the hemolymph to infect the murine liver was severely hindered. These sporozoites also displayed impaired infectivity *in vitro* and motility in glass slides, as well as moderately reduced cell wounding capacity, which could explain their inability to effectively initiate liver infection. Furthermore, we observed that, unlike WT sporozoites, SPATR KD hemolymph sporozoites were incapable of homing to the liver in the first instances following blood dissemination in Kupffer cell-depleted mice, indicating parasite arrest in the sinusoids may have also been compromised. While the defective migratory and infective capacity of SPATR KD hemolymph sporozoites may have been a direct consequence of the downregulation of SPATR, it could have also stemmed from an increase in their permanence time in the vector circulatory system. In general, the *Plasmodium* mutant lines known to likely present a defect in salivary gland invasion also exhibit reduced infectivity and/or motility (Table S3). The fact that all these lines share analogous phenotypes is indicative of one of two options: sporozoites either rely on a diverse roster of proteins whose functions might be intricately related and simultaneously affect a multitude of biological processes, from motility to invasion of the salivary glands and infection of the mammalian host; or impediments to salivary gland invasion induce changes to the sporozoites present in the hemolymph, which result in a decrease in overall fitness and infectivity. These alterations could be intrinsic, meaning that sporozoites may need to timely cross checkpoints to ensure they retain their infective qualities, or exogenous, such as the potential increased contact with antimicrobial peptides and other components of the hemolymph, although we did not detect a decrease of hemolymph sporozoite viability in a SPATR KD line. Nonetheless, the magnitude of the liver infectivity defects varies between the several mutants (Table S3), indicating that the defective phenotype may come from a direct or indirect role of the studied molecule in the process. Indeed, we are studying several *P. berghei* mutant lines whose sporozoites exhibit defective salivary gland invasion and the degree of the liver infectivity deficiency using hemolymph sporozoites varies according to the mutant (data not shown). A milder defective phenotype was obtained when we infected mice with SPATR KD salivary gland sporozoites, as these parasites were capable of infecting mice, unlike their hemolymph counterparts, albeit with a small extension of the prepatent period in comparison to the WT line (0.73 ± 0.57 days, across 7 independent experiments using either SPATR KD clone; Table S2). The minute numbers of sporozoites within the salivary glands made it technically unfeasible to reliably evaluate the motility and infectivity of SPATR KD salivary gland sporozoites *in vitro*, as well as the downregulation of *spatr* transcription in this stage. However, the *in vivo* data indicate that either small amounts of SPATR that may be produced by SPATR KD salivary gland sporozoites are sufficient to secure a certain degree of functionality, or that fully matured sporozoites acquired compensation mechanisms that allowed them to partially revert the phenotype of SPATR KD hemolymph sporozoites, which would be suggestive of functional redundancy. Alternatively, the reduced capacity to invade the salivary glands and, as a consequence, the lower parasite density achieved by these mutant lines in the glands could result in decreased sporozoite fitness. Additionally, it was also shown that saliva components that affect sporozoite functionality are upregulated upon infection (62), which could be dependent on parasite loads. Thus, it is plausible that there is a density threshold within the salivary glands required for the maximization of sporozoite fitness and infectivity. Indeed, SPATR conditional knockout sporozoites generated using the Flp-FRT system were capable of invading the salivary glands and showed no liver stage defects (19). Nevertheless, it is important to note that these recombinase-based systems often fail to achieve full deletion of the target sequence within a population (63, 64), which could be of particular significance in the context of secreted proteins. Moreover, maximum excision of the target locus is only achieved upon a temperature switch, which is usually performed once sporozoites are already present in the salivary glands, as the rise in temperature leads to the blockage of salivary gland colonization (63). Since *spatr* starts being transcribed in oocysts and is then subjected to translational repression (15), this creates a temporal window for sporozoites to produce and store mRNA that can be translated even after the recombination event. Therefore, the study of the importance of genes for the mosquito phase of the life cycle of the parasite using the Flp-FRT system may have additional limitations.

We were unable to generate knockdown lines using the strategy that resorted to the intergenic region upstream of the *cep76* gene, indicating that the promoter elements that drive the expression of this gene in blood stages are not present in the selected genomic region, the levels of *spatr* transcripts were insufficient to sustain parasite survival, or the timing of transcription was not appropriate. On the other hand, the insertion of a second copy of *spatr* under transcriptional regulation of the *hado* promoter in a phenotypically silent locus of chromosome 6 enabled the deletion of the endogenous copy of the gene, originating SPATR KD lines. These lines showed lower levels of *spatr* mRNA throughout sporozoite development and protein levels in hemolymph sporozoites were undetectable by western blot, unlike in SPATR DE parasites. This demonstrated that SPATR is expressed by WT sporozoites prior to salivary gland colonization and that the use of the 2.4 Kb genomic region upstream of the gene that encodes HADO in a promoter exchange system enables the downregulation of highly expressed sporozoite proteins, while sustaining expression in merozoites and, presumably, in ookinetes.

The genetic complementation of a SPATR KD line with an episomally-expressed copy of *spatr* partially reverted the marked reduction of sporozoite numbers in the salivary glands, which confirmed SPATR is required for the invasion of the salivary glands. Remarkably, this infectivity defect of the SPATR KD lines in the mosquito was not accompanied by the expected accumulation of sporozoites in the hemolymph. This could come as a result of the balance between the salivary gland invasion impairment and other factors, such as a potential mild defect in sporozoite egress from oocysts, as suggested by the increased numbers of sporozoites still residing inside oocysts at late time points post-infection and the greater rise in of fully formed SPATR KD sporozoite numbers within oocysts from days 11-13 to day 15, in comparison to the control lines. The similar numbers of hemolymph sporozoites in the first few days after the egress of sporozoites from oocysts is expected to be initiated (day 10 to 11 post-infection) also point towards the attainment of a balance state early on, considering sporozoites residing in the hemolymph did not appear to accumulate in other mosquito tissues and did not exhibit increased mortality.

In conclusion, the loss of SPATR lead to defects in several biological processes necessary for infectivity of hemolymph sporozoites to both the vertebrate and the invertebrate hosts. The function of SPATR in salivary gland colonization and infectivity to the mammalian host might be more crucial in sporozoites that have not yet completed their maturation process (i.e. hemolymph sporozoites) if proteins more highly expressed by salivary gland sporozoites, namely other adhesins, are capable of fulling the same role. This was only made evident upon the early knockdown of *spatr* in the mosquito stages, which also showcases the importance of the selection of the appropriate methods to study discrete aspects of sporozoite biology.

## Materials and methods

### Animals and parasites

Three- to six-week-old C57BL/6, BALB/c, NMRI, CD1 and SWISS mice were purchased from Elevage Janvier, Charles River France or the i3S animal facility. Procedures were conducted in accordance with the guidelines of the Animal Welfare and Ethics Review Body at i3S and the Portuguese National Authorities for Animal Health guidelines, or the Animal Care and Use Committee of Institut Pasteur, following the directive 2010/63/EU of the European Parliament and Council.

All experiments were performed using the *Plasmodium berghei* ANKA strain clone 676cl1 expressing a green fluorescent protein (GFP)-luciferase fusion protein (25). These parasites are herein referred to as wildtype (WT), as they bear the unmodified locus of the gene studied herein. Female *Anopheles stephensi* SDA-500 mosquitoes reared in the Center for the Production and Infection of Anopheles at the Pasteur Institute in Paris, France, were fed on infected SWISS, CD1 or NMRI mice 3–5 days after emergence and kept at 21 ± 0.5 °C and 70% humidity. A feeding using naïve mice was performed 6–7 days after the infectious feeding (26).

Oocyst and salivary gland sporozoites were obtained by mechanical disruption of mosquito midguts or salivary glands, respectively, and collected into Dulbecco’s Phosphate Buffered Saline (DPBS; Gibco). Oocyst sporozoite suspensions were filtered using a 35 µm cell strainer cap (Falcon). To analyze oocysts, midguts of infected mosquitoes were collected at different time points post-infection. To recover hemolymph sporozoites, the haemocoel of mosquitoes was perfused with approximately 15–30 µl of DPBS. The thorax and abdomen of mosquitoes were recovered after the removal of the digestive system, salivary glands and hemolymph. Mosquito material and sporozoites were kept on ice until further processing. Blood schizont purification was performed using a Nycodenz (Axis Shield) gradient following the culture of infected mouse blood in RPMI 1640 (Lonza) supplemented with 50 µg/ml neomycin (Sigma) and 20% heat-inactivated fetal bovine serum (FBS; Biowest) in hypoxic conditions (5% CO_2_, 5% O_2_ and 90% N_2_), for 16 to 18 h at 37 °C and under gentle agitation (27). Free merozoites were isolated via resuspension of mature schizonts using a micropipette or insulin syringe and filtration through a 1.2 µm Acrodisc® filter (Pall Corporation).

### Oligonucleotides

The sequences of all primers (Sigma or Eurofins Genomics) used in this study are shown in Table S1.

### Generation of *P. berghei* transgenic lines

Deletion of the majority of the *spatr* gene (PBANKA_0309500) by double cross-over homologous recombination was attempted using the NotI-linearized PbGEM-270402 vector (*Plasmo*GEM, Sanger) (28, 29). A second construct for total gene deletion was generated by cloning the 5’ (843 bp) and 3’ (292 bp) homology regions amplified by PCR using a Taq DNA polymerase with proofreading activity (Takara) and primers P9 + P12 and P13 + P14 into the pL0001 vector (BEI Resources) via the KpnI/ClaI and EcoRI/BamHI cloning sites, respectively. The linearized constructs were used to transfect synchronized schizonts with the Nucleofector® device (Amaxa). Electroporated parasites were injected intravenously into two mice (parental populations) and positive selection of transfectants was initiated the following day with 70 mg/L pyrimethamine (Sigma) in drinking water. When resistant populations arose, blood was transferred to naïve mice (transfer populations) for a second round of selection with pyrimethamine. Isolation of isogenic lines was performed by limiting dilution *in vivo*, as previously described (27).

To generate SPATR KD lines, we resorted to a two-step strategy: 1) introduction of a second copy of the *spatr* gene under transcriptional control of the intergenic regions upstream of the genes encoding the haloacid dehydrogenase domain ookinete protein (*hado*; PBANKA_0603900) or the putative centrosomal protein 76 (*cep76*; PBANKA_0102600) into a silent intergenic locus in chromosome 6 (30); 2) deletion of the endogenous copy of *spatr*, performed using the PbGEM-270402 vector (*Plasmo*GEM, Sanger). The first step required the insertion of the 5’ (490 bp) and 3’ (474 bp) homology regions amplified by PCR with primer pairs P15 + P16 and P17 + P2 into the KpnI/ClaI and SpeI/SacII cloning sites, respectively, of the pL0001 vector. The genomic region comprised of the open reading frame (1,083 bp) and 396 bp of the downstream sequence were cloned into the construct using the primer pair P18 + P19 and the BamHI and SpeI restriction enzymes. The 5’ intergenic regions of *hado* (2,414 bp) and the putative *cep76* (845 bp) were amplified with primer pairs P20 + P21 and P22 + P23, respectively, and inserted into the plasmid using the EcoRV/BamHI restriction sites. Transfections were performed as before, and selection of transfectants was achieved using pyrimethamine as previously described (step 1) or 10–16 mg/kg/day of WR99210 (Jacobus Pharmaceutical Company) injected subcutaneously for 3 consecutive days (step 2). Parasites were then cloned by limiting dilution. Two isogenic lines (SPATR KD^V1^ and SPATR KD^S16^) were used in this study.

To generate a parasite line expressing a c-Myc-tagged version of SPATR, the WT *spatr* open reading frame was replaced by a copy of *spatr* fused with a sequence encoding a 2x c-Myc tag at the C-terminus, by single cross-over homologous recombination. Since the linearization of the transfection vector was restricted to the AdhI restriction site within the *spatr* gene, modifications were made to pL0033 (BEI Resources) in in order to remove the AdhI restriction site originally present in the plasmid. The pL0033 fragment containing the ampicillin resistance cassette and the AdhI site that resulted from SapI and NcoI digestion was swapped by the fragment containing the kanamycin marker obtained by digestion of pET-28a(+) (Novagen) with the same restriction enzymes, generating pL0033_Kan. The genomic region encompassing 439 bp of the 5’ intergenic region and the open reading frame of *spatr* excluding the termination codon was then amplified by PCR using primers P24 + P25 and cloned into pL0033_Kan using the NotI and NcoI restriction sites. The 3’ regulatory sequence of *spatr* (358 bp) amplified with P26 + P27 was inserted downstream of the sequence encoding the 2x c-Myc tag using the FseI/NdeI restriction sites. A point mutation was inserted in the start codon to prevent the translation of the full-length untagged SPATR due to the duplication of part of the locus resulting from the genetic recombination event. To do this, the region containing the first 13 nucleotides of the coding sequence and the entire 5’ intergenic region previously inserted in the construct was swapped for the sequence amplified using primers P24 + P28 and using the NotI/AhdI restriction sites. Parasites were transfected using the AhdI-linearized construct. Transfectant selection with pyrimethamine and cloning was performed as described above.

To genetically complement the SPATR KD^V1^ line, we transfected parasites with constructs generated by modifications of a plasmid containing the centromere sequence of the *P. berghei* chromosome 5 (31–33). The sequence comprised of the open reading frame of *spatr* and the 3’ intergenic region (1491 bp) was amplified with primers P29 + P30 and cloned into the episome using SacII and XhoI. The gene was put under transcriptional control of the 5’ intergenic region of *spatr* (1,000 bp) or the promoter of the gene encoding the thrombospondin related anonymous protein (*trap*; PBANKA_1349800; 1,501 bp) (23), which were amplified using primer pairs P31 + P32 or P33 + P34, respectively, and inserted into the PstI/SacII-digested transfection vector. This plasmid also contained the promoter of the eukaryotic elongation factor 1A gene (*eef1αa*; PBANKA_1133400) (34) constitutively driving the expression of the RedStar reporter. This allowed for the enrichment of episome-carrying blood stages by fluorescence-activated cell sorting (FACS) using a FACS Aria II machine (Benton Dickson) (25). Blood was diluted in DPBS containing 2% FBS, and 30 to 1×10^4^ GFP-positive/RFP-poritive erythrocytes were sorted at room temperature, collected in 100-150 µl of DPBS containing 10% FBS and immediately injected intravenously in recipient mice. Several successive enrichment steps were performed to increase the percentage of RFP+ parasites within the populations. FACS data was analyzed using the FlowJo software version 10.

All PCR products were primarily cloned into the pGEM-T Easy vector (Promega) and sequenced (LightRun, Eurofins Genomics) before being subcloned into the respective transfection vector. All restriction enzymes used in this study were from New England Biolabs, with the exception of AdhI (Fermentas).

### Genotyping of transgenic parasites

Blood collected from infected animals was filtered with a Plasmodipur leukocyte filter (EuroProxima) and erythrocytes were lysed with saponin (Fluka) at 0.15%. Genomic DNA was extracted and purified from the resulting parasite pellet using the QIAamp DNA Blood Mini Kit (Qiagen). Integration of the constructs in the correct loci and absence of the WT loci were then confirmed by PCR using the Phusion DNA Polymerase (New England Biolabs).

For Southern blot, 5 µg of genomic DNA were digested with EcoRI and EcoRV, separated on a 0.8% agarose (NZYTech) gel and transferred to a Hybond-N^+^ nylon membrane (Amersham). The intergenic sequence at 3’ of *spatr* amplified using primer pair P13 + P14 was used as a probe. Labelling of the probe and signal generation and detection were performed with the AlkPhos Direct Labeling and Detection System with CDP-Star chemiluminescent detection reagent (Amersham). Signal detection was performed using a ChemiDoc Imaging System (Bio-Rad).

### Analysis of blood stages

The growth of asexual blood stages was assessed by infecting NMRI mice at a parasitaemia of 0.01% and counting infected red blood cells in blood smears stained with Hemacolor (Merck). The percentage of total gametocytes was also determined by counting in blood smears. To assess microgamete exflagellation, a drop of blood was collected from an infected mouse and placed on a glass slide. Exflagellation was induced by breathing on a coverslip, placing it on top of the drop of blood and incubating for 10 min at room temperature. Exflagellation centers were counted in 5 fields using a widefield microscope at 40X magnification.

### Reverse transcriptase quantitative PCR (RT-qPCR)

Total RNA from merozoites, oocysts and sporozoites was extracted using TRIzol LS Reagent (Invitrogen) and Phase Lock Gel Heavy tubes (5PRIME), followed by the removal of contaminating DNA with TURBO DNase (Invitrogen). Superscript® II Reverse Transcriptase (Invitrogen) was used to synthesize cDNA and the determination of *spatr* (primers P35 + P36), *hado* (primers P37 + P38), *cep76* (primers 39 + 40) and trap (primers 41 + 42) transcript levels by RT-qPCR was performed using iQ or iTaq™ Universal SYBR Green supermix (Bio-Rad) and the iQ5 or CFX96 real-time PCR detection systems (Bio-Rad). We used *hsp70* as the reference gene (primers P43 + P44) and the 2^-Δ*CT*^ method to determine the relative gene expression. Quantification of sporozoites associated with the abdomen and thorax following the removal of the digestive tracts, salivary glands and hemolymph was performed on day 15 post-infection by detection of 18S ribosomal RNA (primers P45 + P46) and interpolation from a standard curve constructed with salivary gland sporozoites processed the same way as the mosquito carcasses. Amplification efficiencies were between 90 and 110%.

### Analysis of oocyst development

Mosquito midguts were collected from day 6 to day 15 post-infection. To calculate oocyst number and size, midguts were collected into an 18-well µ-slide (ibidi) containing DPBS and images were taken using a Axio Observer Z1 fluorescence microscope (Zeiss) at 5X magnification. From day 11 post-infection, midguts were crushed, and the hemolymph and salivary glands were collected for sporozoite quantification in all compartments.

### Sporozoite viability

Hemolymph sporozoites collected on day 21 post-infection were diluted in DPBS containing propidium iodide (PI; Invitrogen) at a final concentration of 5 µg/ml, transferred to an 18-well µ-slide (ibidi), and centrifuged for 5 min at room temperature and 500 x g. Photos were taken using an IN Cell Analyzer 2000 (GE Healthcare) and PI incorporation was evaluated with the ImageJ software.

### Production of recombinant protein

The coding sequence of *spatr* lacking the 22 amino acids of the predicted signal peptide (SignalP-4.1 server) was amplified from salivary gland sporozoite cDNA using primer pair P47 + P48 and cloned in frame into the expression plasmid pET-28a(+) using NcoI and XhoI restriction sites. A recombinant SPATR (rSPATR) fused to a polyhistidine tag (His-tag) was produced in BL21-CodonPlus*-*RIL competent cells (Invitrogen) and protein purification was performed under denaturing conditions using Ni-NTA Agarose (Qiagen) following the instructions of the manufacturer. Desalting and concentration of rSPATR was performed using a Vivaspin 20 3,000 MWCO PES (Sartorius) centrifugal concentrator and DPBS. To remove endotoxins, the recombinant protein was briefly vortexed in the presence of 1% Triton X-114 (Sigma), incubated on ice for 15 min, vortexed again, and incubated at 37°C for 10 min (35). The mixture was centrifuged at room temperature and at 2,000 x g for 10 min and the top phase was collected. The procedure was performed a total of 3 times and endotoxin levels were determined using the Pierce™ LAL Chromogenic Endotoxin Quantitation Kit (Thermo Scientific).

### Production of polyclonal antibodies

A mouse polyclonal antibody was generated by immunizing 6-week-old BALB/c mice intraperitoneally 3 times with 25 µg of rSPATR adjuvanted with 50 µg of Poly(I:C) (polyinosinic-polycytidylic acid) as an adjuvant, followed by a fourth booster dose of 50 µg of rSPATR and 50 µg of Poly(I:C). All doses were administered 2 weeks apart. Serum was collected 12 days after the final dose.

### SDS-Page and Western blot

Purified merozoites and sporozoites were lysed in Laemmli sample buffer at 95°C for 10 min. Sample volumes containing 1.5-4×10^6^ merozoites, 1-4×10^5^ sporozoites or 100 to 5,000 ng of rSPATR were separated on a 12% acrylamide gel by SDS-PAGE. SDS-PAGE gels were stained with Coomassie blue and scanned using a GS-800 densitometer (Bio-Rad). For Western blot, the transfer to a PVDF membrane was performed with a Trans-blot Turbo Transfer System (Bio-Rad). The membranes were blocked using DPBS with 0.1% Tween 20 (Sigma) and 5% skim milk for at least 1 h at room temperature. Incubations with primary mouse α-rSPATR polyclonal antibodies (1:200 dilution), rabbit α-His-tag antibodies (0.1 µg/ml; MicroMol), mouse α-c-Myc antibodies (2.5 to 5 µg/ml; Merck), or mouse α-CSP 3D11 monoclonal antibodies (6, 36) (0.3 µg/ml; BEI resources), diluted in the blocking solution, were performed overnight at 4°C (α-rSPATR) or for 1 h at room temperature (α-His-tag and α-CSP 3D11). The membranes were washed and incubated with the secondary HRP-conjugated anti-rabbit (1:10,000) or anti-mouse (1:5,000; SouthernBiotech) antibodies for 1 h at room temperature. Signal detection was performed using SuperSignal West Pico Chemiluminescent Substrate (Thermo Scientific) and a ChemiDoc Imaging System (Bio-Rad), or with Amersham Hyperfilm ECL (cytiva) and a FPM-100A film processer (Fujifilm).

### Immunofluorescence assays and fluorescence microscopy of live sporozoites

Hemolymph sporozoites collected on days 19 to 24 post-infection were centrifuged for 5 min at 450 x g and 4 °C in an 8-well Lab-Tek chamber slide (Nunc) or a 384-well plate (Greiner), fixed with 4% formaldehyde (Merck), washed 5 times with DPBS, and blocked with DPBS containing 5% FBS for at least 30 min at room temperature. Sporozoite activation, when applicable, was achieved by incubation for 10 min at 37 °C in DMEM containing 5% FBS, prior to fixation. Incubation with α-CSP 3D11 mouse monoclonal antibodies (2.3 µg/ml) was performed for 1 h at room temperature. Incubation with α-TRAP rabbit polyclonal antibodies (1:10,000) was performed overnight at 4 °C. The secondary antibodies, either goat α-mouse or α-rabbit Alexa Fluor 568 antibodies (2 µg/ml; Invitrogen), were added and incubation for 30 min at room temperature ensued. Finally, nuclei were stained with DAPI (1 µg/ml; Acros Organics) for 10 min at room temperature and the slides mounted with a 90% glycerol (Alfa Aesar), 0.5% (w/v) n-propyl gallate (Sigma), 20 mM Tris-HCl (Sigma) pH 8.0 antifade solution. All stainings were performed using antibodies diluted in the blocking solution and were followed by 5 washes with DPBS. Live sporozoites were centrifuged for 3 min at 450 x g in a 18-well µ-slide (ibidi), at room temperature. Sporozoites were then analyzed using a DMI6000 FFW microscope (Leica Microsystems) and the ImageJ software.

### Transmission electron microscopy

For the ultrastructure analysis, salivary glands collected from infected mosquitos at day 27 post-infection were fixed in a solution of 2,5% glutaraldehyde (Electron Microscopy sciences) with 2% formaldehyde (Electron Microscopy sciences) in 0.1 M sodium cacodylate buffer (pH 7.4) for 24 h, at room temperature. After washing in buffer and two hours in post-fixating 2% osmium tetroxide (Electron Microscopy sciences) in 0.1M sodium cacodylate (Electron Microscopy sciences) buffer (pH 7.4), tissues were washed in water and then incubated with 1% uranyl acetate (Electron Microscopy sciences) overnight. Samples were dehydrated through graded series of ethanol, and embedded in EPON resin (Electron Microscopy sciences). Ultra-thin sections (50 nm thickness) were cut on a RMC Ultramicrotome (PowerTome) using Diatome diamond knifes, mounted on mesh copper grids (Electron Microscopy Sciences), and stained with uranyl acetate substitute (Electron Microscopy Sciences) and lead citrate (Electron Microscopy Sciences) for 5 min each. Samples were viewed on a JEM 1400 transmission electron microscope (JEOL) and images were digitally recorded using an 1100W CCD digital camera (ORIUS).

### *In vitro* invasion and development assays

Between 7×10^4^ and 1.2×10^5^ HepG2 cells (ATCC) per well were seeded in an 8-well Lab-Tek chamber slide (Nunc), precoated with rat tail collagen type I (Sigma) at 200 µg/ml and incubated at 37°C overnight in DMEM (Gibco or Lonza) supplemented with 10% FBS, in a humidified incubator with 5% CO_2_. Hemolymph from mosquitos containing oocysts was collected by perfusion with DPBS at 4°C on days 17 to 21 post-infection. Sporozoites were counted, diluted in an equal volume of DMEM containing 20% FBS and 200 U/ml of penicillin/streptomycin (Lonza), and ∼1.5×10^4^ hemolymph sporozoites were added per well. The slides were centrifuged at 450 x g for 5 min at room temperature to promote contact between sporozoites and HepG2 cells, and incubated at 37 °C. To evaluate the capacity of sporozoites to infect host cells, the preparations were fixed with 4% formaldehyde (Merck) 1.5–2 h after the addition of the sporozoites. The medium in slides prepared to assess EEF development was changed daily until the fixation time point (24 or 48 h). Intracellular and extracellular sporozoites were discriminated via an anti-CSP double staining. Preparations were blocked with DPBS containing 5% FBS for 30 min at room temperature, incubated with α-CSP 3D11 mouse monoclonal antibodies (2.3 µg/ml) for 1 h at room temperature, washed with DPBS, incubated with goat α-mouse Alexa Fluor 568 antibodies (2 µg/ml; Invitrogen) for 1 h at room temperature, and washed again. The incubation with the primary α-CSP antibodies was repeated after permeabilization with 1% Triton ×100 (Sigma) and washing. After the washing steps, incubation with goat α-mouse Alexa Fluor 488 antibodies (2 µg/ml; Invitrogen) was performed for 1 h at room temperature. Nuclei were then stained with DAPI diluted 1:5,000 in DPBS for 10 min at room temperature. To facilitate the detection of weakly-fluorescent liver stages, cells were permeabilized as described above and sequentially stained with primary rabbit α-GFP antibodies (1:250; MBL) and secondary goat α-rabbit Alexa Fluor 488 antibodies (2 µg/ml; Invitrogen) for 1h at room temperature. Finally, the wells were washed and the slides mounted with antifade solution. Image acquisition and analysis was performed as previously described (26). Briefly, 25 images per well, corresponding to a total area of 0.57 mm^2^, were taken using IN Cell Analyzer 2000 (GE Healthcare). HepG2 nuclei as well intracellular and extracellular sporozoites were counted using the IN Cell Developer Toolbox v.1.9.2 software (GE Healthcare). The area of liver stages was also determined using this software.

### *In vitro* cell wounding assay

Cell wounding assays were performed using a previously described protocol (37), with modifications. Briefly, HepG2 cells (ATCC) were seeded at 2×10^4^ cells per well on a collagen-coated 96-well plate (Orange Scientific) and incubated at 37 °C overnight in DMEM containing 10% FBS. Hemolymph and salivary gland sporozoites collected in DPBS on day 19 post-infection were diluted in an equal volume of DMEM containing 20% FBS, 200 U/ml of penicillin/streptomycin and PI at 10 µg/ml, and 2.7×10^4^ to 4×10^4^ sporozoites were added per well. Controls without PI and/or parasites were also included. The plate was centrifuged at 500 x g for 5 min at room temperature and incubated for 1 h at 37 °C in a humidified incubator with 5% CO_2_. The medium was removed, and the cells were washed and digested with trypsin (Gibco) for 10 min at 37 °C. DMEM supplemented with 10% FBS was added, cells were centrifuged at 500 x g for 5 min at room temperature and resuspended in DPBS containing 2% FBS. The percentage of PI-positive cells was determined by flow cytometry analysis using a FACSCanto II (BD Biosciences) and FlowJo software version 10.

### *In vivo* infections with sporozoites

To evaluate the infectivity of the mutant lines, 1.1×10^4^ to 6×10^4^ hemolymph or 200 to 2,500 salivary gland sporozoites collected on days 18 to 24 or 19 to 27 post-infectious blood meal, respectively, were injected intravenously into C57BL/6 mice. Parasite burdens in the liver were evaluated by bioluminescence imaging at day 1 and/or 2 post-infection. Isoflurane-anaesthetized mice were shaved and injected subcutaneously with 2.4 mg of D-luciferin potassium salt (Perkin Elmer) dissolved in DPBS. Mice were transferred to the IVIS Lumina LT system (Perkin Elmer) stage and kept under anesthesia during signal acquisition, which was initiated 5 min after the administration of the substrate. Parasitemias were calculated from day 3 post-infection by counting infected red blood cells in blood smears stained with Hemacolor (Merck).

The targeting of the liver by hemolymph sporozoites in the first minutes post-infection was assessed using whole-mouse bioluminescence imaging (6). Mice were pretreated with 10 μl/g of clodronate liposomes (Liposoma BV) 2 to 4 days prior to infection to deplete Kupffer cells (4). C57BL/6 mice were previously shaved and infected with 9×10^4^ to 1.5×10^5^ hemolymph sporozoites by intravenous injection and anesthetized, and D-luciferin was administered 4 min after infection. Signal detection was initiated 7 min post-infection and performed for 5 min. Living Image software (Perkin Elmer) was used to control que acquisition process, to quantify the bioluminescent signal in regions of interest (ROIs) and to generate the photograph and signal map overlays.

### Gliding assays

The evaluation of sporozoite motility *in vitro* was performed by live microscopy on 18-well µ-slides (ibidi). Sporozoites were collected from the hemolymph or the salivary glands of infected mosquitoes on days 18–19 or 24–25 post-infectious feeding, respectively. Salivary glands were mechanically disrupted using a pestle and filtered using a 35 µm cell strainer cap (Falcon). Sporozoite suspensions prepared in DPBS and kept on ice were diluted in equal parts of DMEM (4.5 g/l glucose) with 10% FBS and transferred to the slide at a final concentration of approximately 5,000 parasites per well. Centrifugation at 400 x g, 4 °C, for 3 min ensued, to allow the parasites to attach to the bottom of the well. The slide was then transferred to the inverted phase contrast microscope (Axio Observer Z1) and incubated at 37°C, 5% CO_2_, for 3 min. Up to 3 videos of 3 min were sequentially recorded in different fields of each well using the 20X objective, and the trajectories and speed of individual parasites were analyzed using the ImageJ software. Motile behaviors were classified as follows: motile ≥ 1 circle – motile sporozoites capable of completing at least a full circle; motile < 1 – motile sporozoites that did not complete a full circle; waving – sporozoites that at any point attached to the bottom of the well, often through one of the poles, without displacement; drifting – parasites that did not at any point attach to the bottom of the well; immotile – sporozoites that remained stationary during the duration of the video.

### Statistical analysis

Statistical analyses were performed using the GraphPad Prism Software (version 8.0). Statistical significance: *p* < 0.05 (*), *p* < 0.01 (**), *p* < 0.001 (***), *p* < 0.0001 (****).

## Acknowledgements

We would like to thank Prof. Anabela Cordeiro da Silva from the IBMC/i3S for the hosting conditions. We would like to thank the Wellcome Sanger Institute for providing the *Plasmo*GEM construct PbGEM-270402 and Daniel Bargieri from the University of São Paulo, Brazil for providing the RFP-CEN5 plasmid. The following reagents were obtained through BEI Resources, NIAID, NIH: (a) Plasmid pL0001, for Transfection in *P. berghei*, MRA-770, contributed by Andrew P. Waters; (b) Plasmid pL0033, for Transfection in Plasmodium berghei, MRA-802, contributed by Andrew P. Waters; (c) Hybridoma 3D11 Anti-Plasmodium berghei 44-Kilodalton Sporozoite Surface Protein (Pb44), MRA-100, contributed by Victor Nussenzweig.

This work was supported by funds from project Norte-01-0145-FEDER-000012 - Structured program on bioengineered therapies for infectious diseases and tissue regeneration, supported by Norte Portugal Regional Operational Programme (NORTE 2020), under the PORTUGAL 2020 Partnership Agreement, through FEDER. This work also received funds from the Fundação para a Ciência e Tecnologia (FCT)/Ministério da Ciência, Tecnologia e Ensino Superior (MECTES) co-funded by FEDER (PTDC/SAU-PAR/31340/2017) under the Partnership agreement PT2020, through the Research Unit No. 4293. J.T. is an Investigator funded by National funds through FCT and co-funded through European Social Fund within the Human Potential Operating Programme (CEECIND/02362/2017). D.M.C., M.S. and A.R.T. are funded by FCT individual fellowships SFRH/BD/123734/2016, SFRH/BD/133485/2017 and SFRH/BD/133276/2017 respectively). We also acknowledge the support of the i3S Advanced Light Microscopy and Histology and Electron Microscopy Scientific Platforms, members of the PPBI (PPBI-POCI-01-0145-FEDER-022122).

## Supplementary figure legends

**Fig S1.**
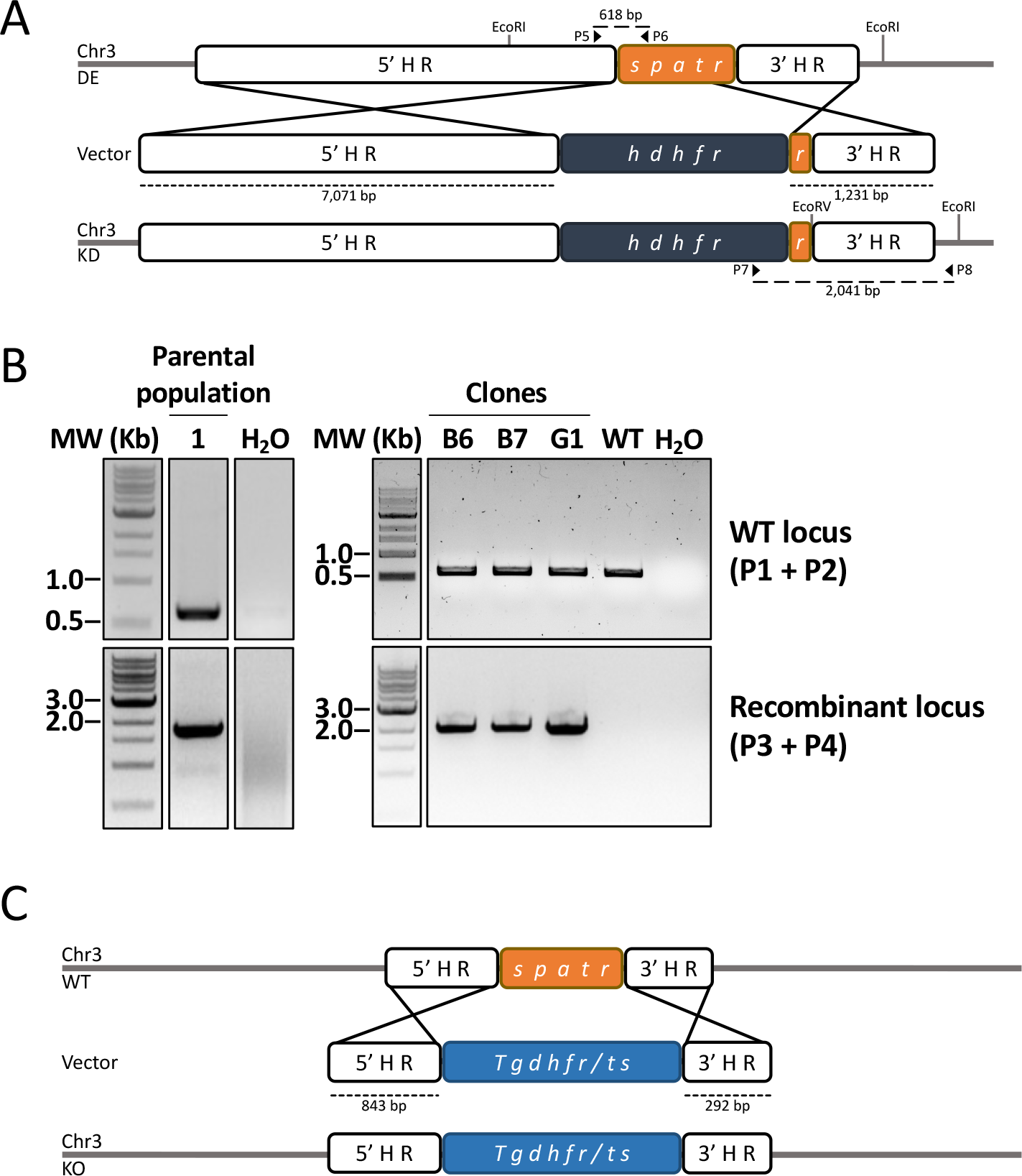
Attempts to knockout *spatr* by targeted gene deletion. (A) Schematic representation of the gene deletion strategy using the PbGEM-270402 vector. The sizes of homology regions are shown next to dashed lines. Sequences are not to scale. (B) Genetic analysis of transfected populations by PCR. The primers used for the genetic analysis and the amplicon sizes (long dashes) are depicted in panel A. (C) Schematic representation of the alternative knockout strategy using the modified pL0001 vector. The sizes of homology regions are shown next to dashed lines. Sequences are not to scale. bp: base pairs. Chr: chromosome. *hdhfr*: human dihydrofolate reductase. HR: homology region. Kb: kilobases. KO: knockout. MW: molecular weight. *Tgdhfr/ts*: *Toxoplasma gondii* dihydrofolate reductase-thymidylate synthase.

**Fig S2.**
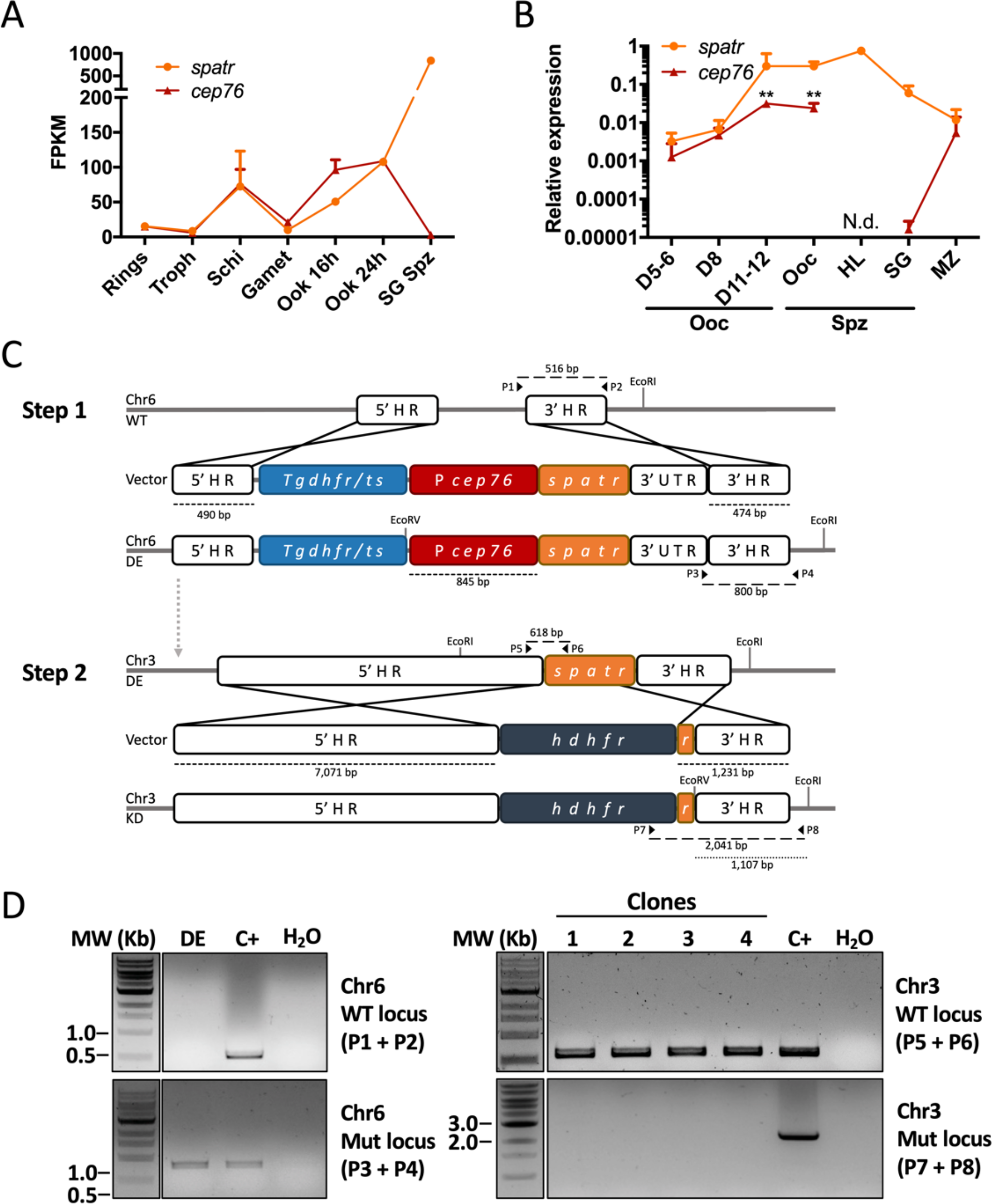
Promoter swap using the intergenic region upstream of the putative *cep76*. (A) RNA-seq data of the putative *cep76* gene in different stages of the *P. berghei* life cycle. Data from published datasets [REFS]. (B) Expression of the putative *cep76* gene relative to *hsp70* in different stages and days post-infection, assessed by RT-qPCR using independently prepared RNA samples. The *spatr* transcription data shown in Fig. 1A and 1B were plotted in panels A and B, respectively, to facilitate comparison the of the transcriptional profiles. Mean values + standard deviation are represented in both panels and statistical analysis in panel B was performed using the Kruskal-Wallis test with Dunn’s multiple comparisons test. Non-significance is not represented. (C) Stepwise schematic representation of the promoter exchange strategy using the promoter of the putative *cep76* gene. The sizes of genomic regions are shown next to dashed line. Sequences are not to scale. (D) Genetic analysis of transfected populations by PCR. Primers used for the genotyping and amplicon sizes are depicted in panel C. bp: base pairs. C+: positive control (WT or transfer population on the top or bottom gels, respectively). Chr: chromosome. DE: double-expressor line (intermediate line that differs from the SPATR DE line originating from the *hado* promoter strategy used as a control in the remaining experiments). Gamet: gametocytes. FPKM: fragments per kilobase of transcript per million mapped reads. *hdhfr*: human dihydrofolate reductase. HL: hemolymph. HR: homology region. Kb: kilobases. KD: knockdown (expected final line that could not be isolated). Mut: mutant. MW: molecular weight. Ooc: oocyst(s). Ook: ookinetes. Schi: schizonts. SG: salivary gland. Spz: sporozoites. *Tgdhfr/ts*: *Toxoplasma gondii* dihydrofolate reductase-thymidylate synthase. Troph: trophozoites. UTR: untranslated region.

**Fig S3.**
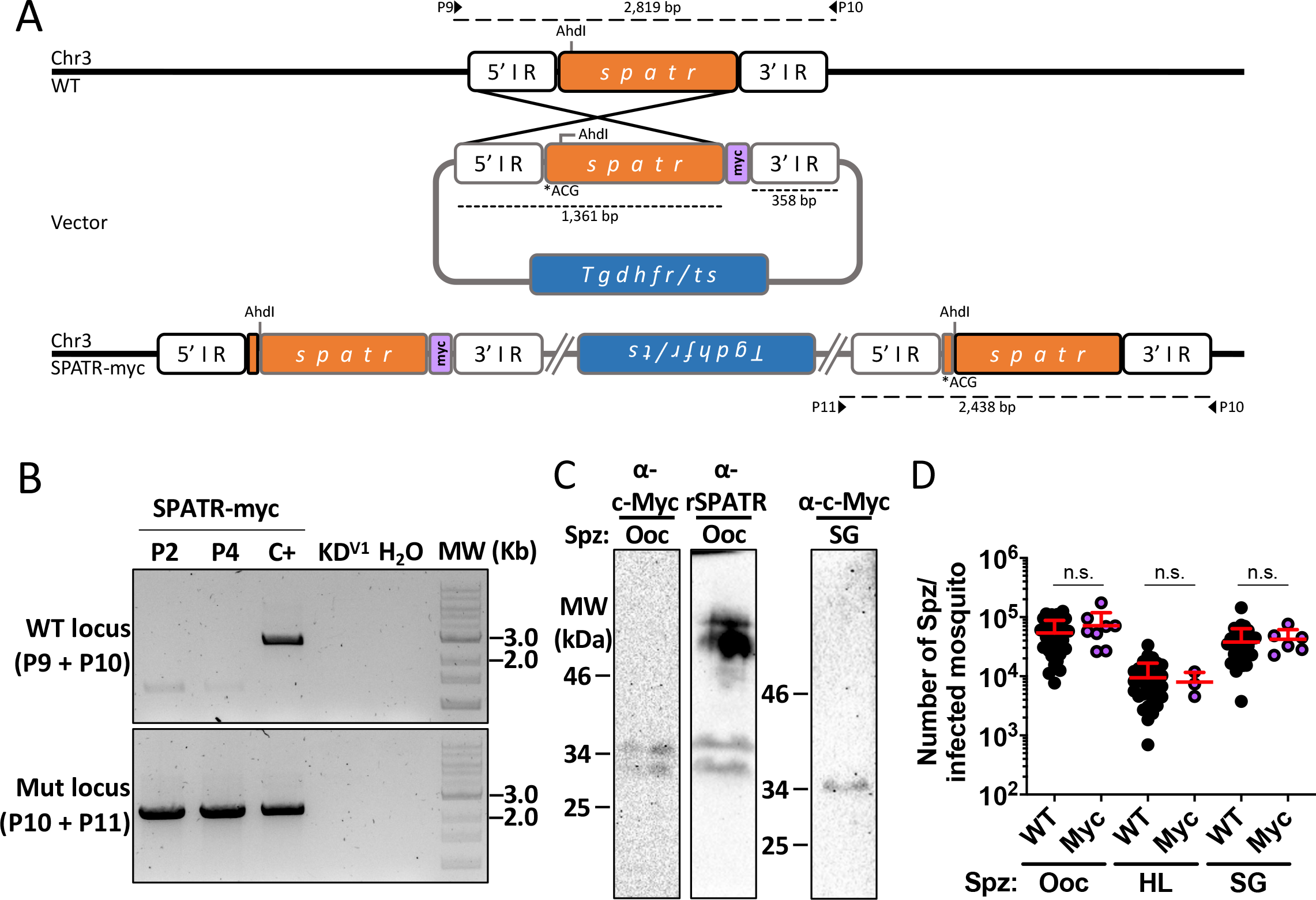
Generation of a line expressing a c-myc-tagged SPATR. (A) Schematic representation of the generation of the SPATR-myc line, which expressed SPATR fused to two c-myc sequences (myc) at C-terminal. The duplicated copy of the gene at 3’ contains a mutated start codon (ACG) to prevent the expression of the full length and untagged SPATR. The sizes of the homology regions are shown below dashed lines, under the vector scheme. Sequences are not to scale. (B) Genetic analysis of two clonal populations by PCR. A transfer population (TP) and SPATR KD^V1^ were used as positive and negative controls, respectively. The primers used for the genetic analysis and the amplicon sizes are depicted in panel A. (C) Reactivity of α-c-Myc and α-rSPATR antibodies to extracts prepared with 4×10^5^ SPATR-myc oocyst sporozoites or 1.5×10^5^ SPATR-myc salivary gland sporozoites, evaluated by western blot. (D) Numbers of SPATR-myc (Myc) oocyst, hemolymph and salivary gland sporozoites per mosquito. WT values are also shown in Fig. 2E. Mean values + standard deviation are represented and statistical analysis was performed using the Mann-Whitney test. n.s.: non-significant. bp: base pairs. C+: positive control (WT or transfer population on the top or bottom gels, respectively). Chr3: chromosome 3. HL: hemolymph sporozoites. IR: intergenic region. Mut: mutant. MW: molecular weight. SPATR My Ooc: oocyst. SG: salivary gland. Spz: sporozoites. *Tgdhfr/ts*: *Toxoplasma gondii* dihydrofolate reductase-thymidylate synthase.

**Fig S4.**
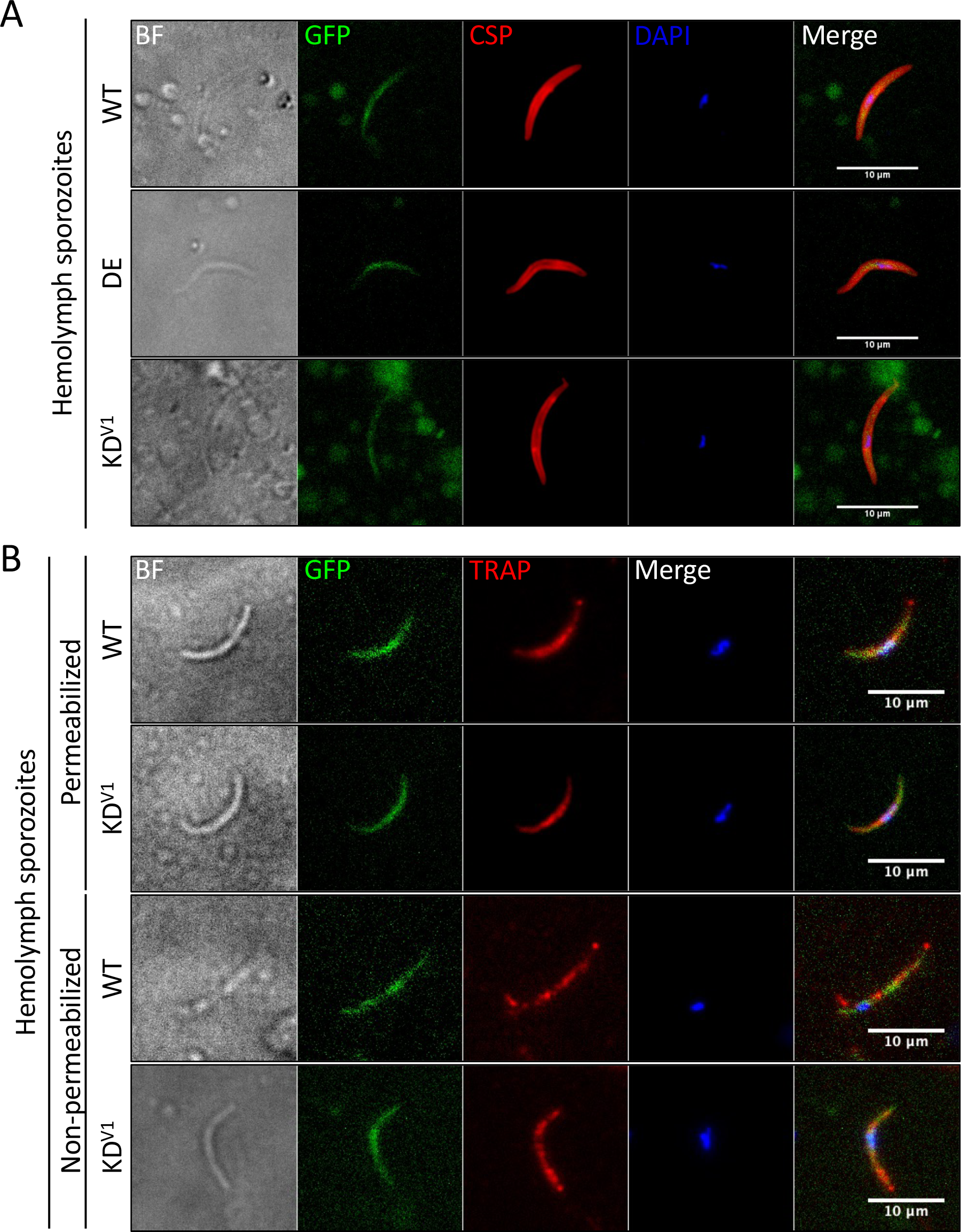
Morphology of KD sporozoites and expression of surface molecules. **(A, B)** Expression and localization of CSP (A) and TRAP (B) in non-activated and activated hemolymph sporozoites, respectively, evaluated by fluorescence microscopy. BF: brightfield. DE: SPATR double-expressor. KD^V1^: SPATR knockdown V1 clone.

**Fig S5.**
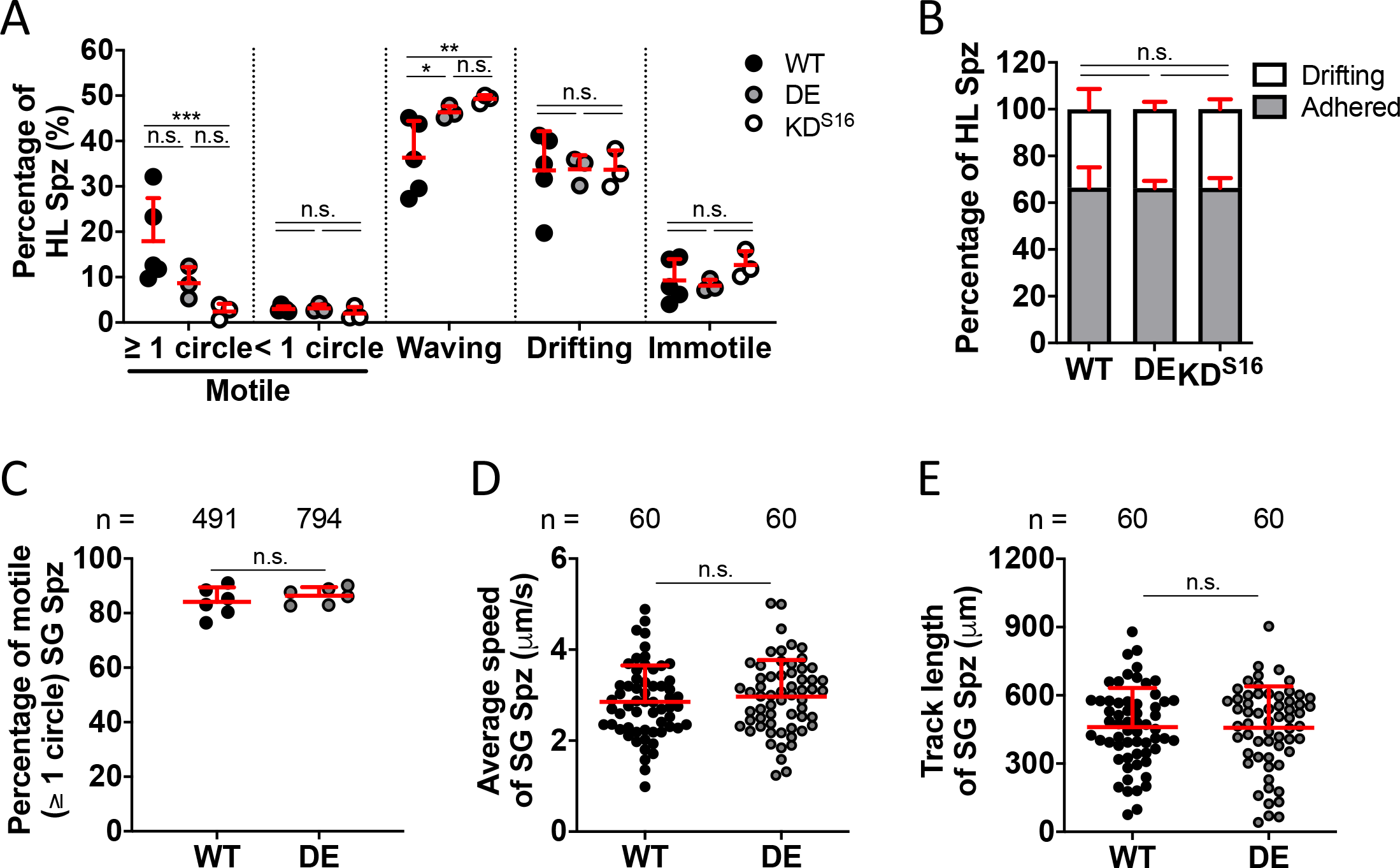
Motile behaviors of SPATR KD and SPATR DE sporozoites. (A) Percentage of hemolymph sporozoites exhibiting distinct motility-associated behaviors. Each symbol represents the mean value of an independent experiment and a total of at least 1223 sporozoites per line was analyzed. (B) Percentage of drifting or adhered hemolymph sporozoites. The percentage of adhered sporozoites was calculated by adding the percentages of motile, waving and immotile parasites in each experiment. Mean values + standard deviation are represented in both panels. (C–E) Percentage of motile salivary gland sporozoites (C), and their average speed (D) and track length (E). Numbers above the plots indicate the total number of sporozoites analyzed. The symbols in panel C represent the percentages obtained in different videos, rather than individual sporozoites analyzed. Statistical analyses were performed using the Kruskal-Wallis with Dunn’s multiple comparisons test (A–B) or the Mann-Whitney test (C–E). n.s.: non-significant. DE: SPATR double-expressor. HL: hemolymph. KD^S16^: SPATR knockdown S16 clone. SG: salivary gland. Spz: sporozoites.

## Supplementary table legends

**Table S1.**
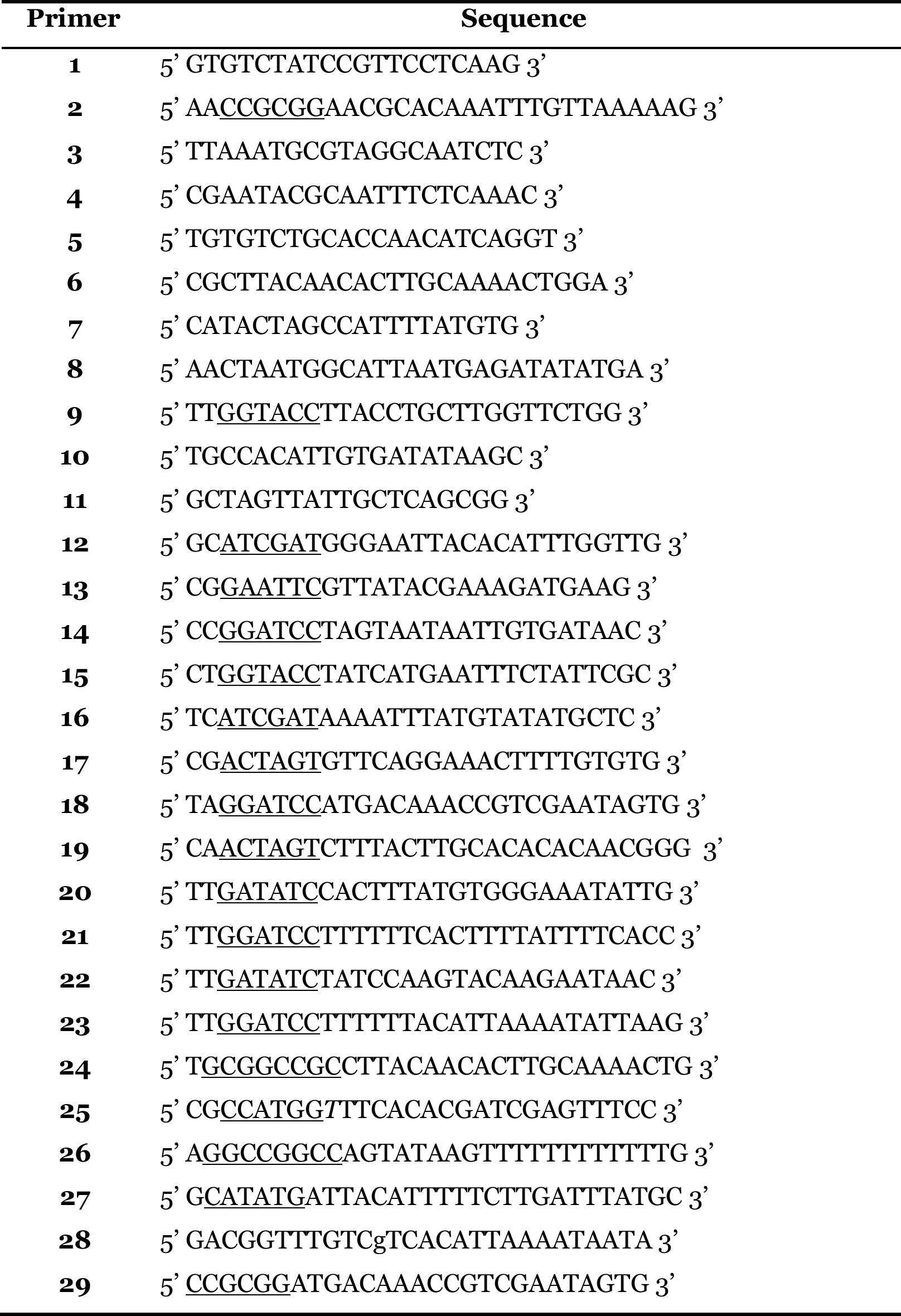
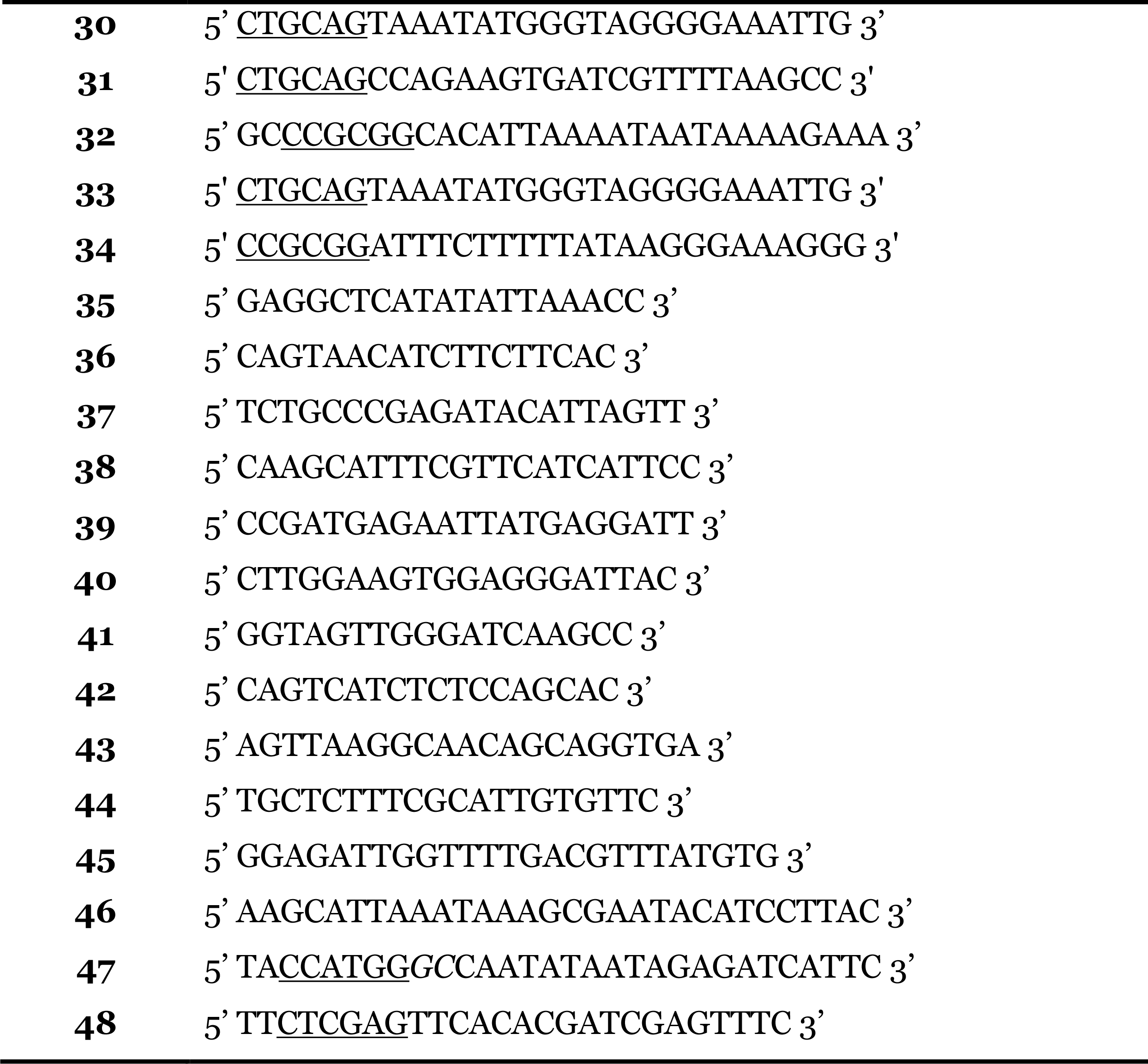
Oligonucleotides. Restriction sites are underlined. Bases in italic were introduced to ensure in-frame cloning. Lower case letters represent point mutations.

**Table S2.** Initiation of blood stage infection by hemolymph and salivary gland sporozoites. The average prepatency delay versus mice infected with WT salivary gland (SG) or hemolymph (HL) sporozoites (Spz) in independent infections carried out with inocula. SG Spz inocula ranged from 200 to 2,500 SG sporozoites is shown. ^†^: Infections with WT and KD lines were performed with 15,000 to 60,000 HL Spz. Only one mouse injected with 20,000 presented parasitemia. ^#^: one mouse was injected with 11,000 hemolymph sporozoites (HL Spz) and did not show parasitemia and the two infected mice were injected with 60,000 HL Spz. No infections with WT were performed simultaneously. COMP^Ptrap^: SPATR COMP^Ptrap^. DE: SPATR DE. n.a.: not applicable. KD^V1/S16^: SPATR KD^V1/S16^.

| Line | SG Spz |  | HL Spz |  |
| --- | --- | --- | --- | --- |
|  | Prepatency<br>increase (days) | % of infected<br>mice | Prepatency<br>increase (days) | % of infected<br>mice |
| WT | n.a. | 27/29 (93%) | n.a. | 16/16 (100%) |
| DE | 0.44 ± 0.10 | 11/12 (92%) | 0.32 ± 0.34 | 11/13 (85%) |
| KD<sup>V1/S16</sup> | 0.73 ± 0.57 | 18/21 (86%) | 1<sup>†</sup> | 1/24 (4%)<sup>†</sup> |
| COMP<sup>Ptrap</sup> | 0.34 ± 0.57 | 6/7 (86%) | n.a.<sup>#</sup> | 2/3 (67%)<sup>#</sup> |

**Table S3.**
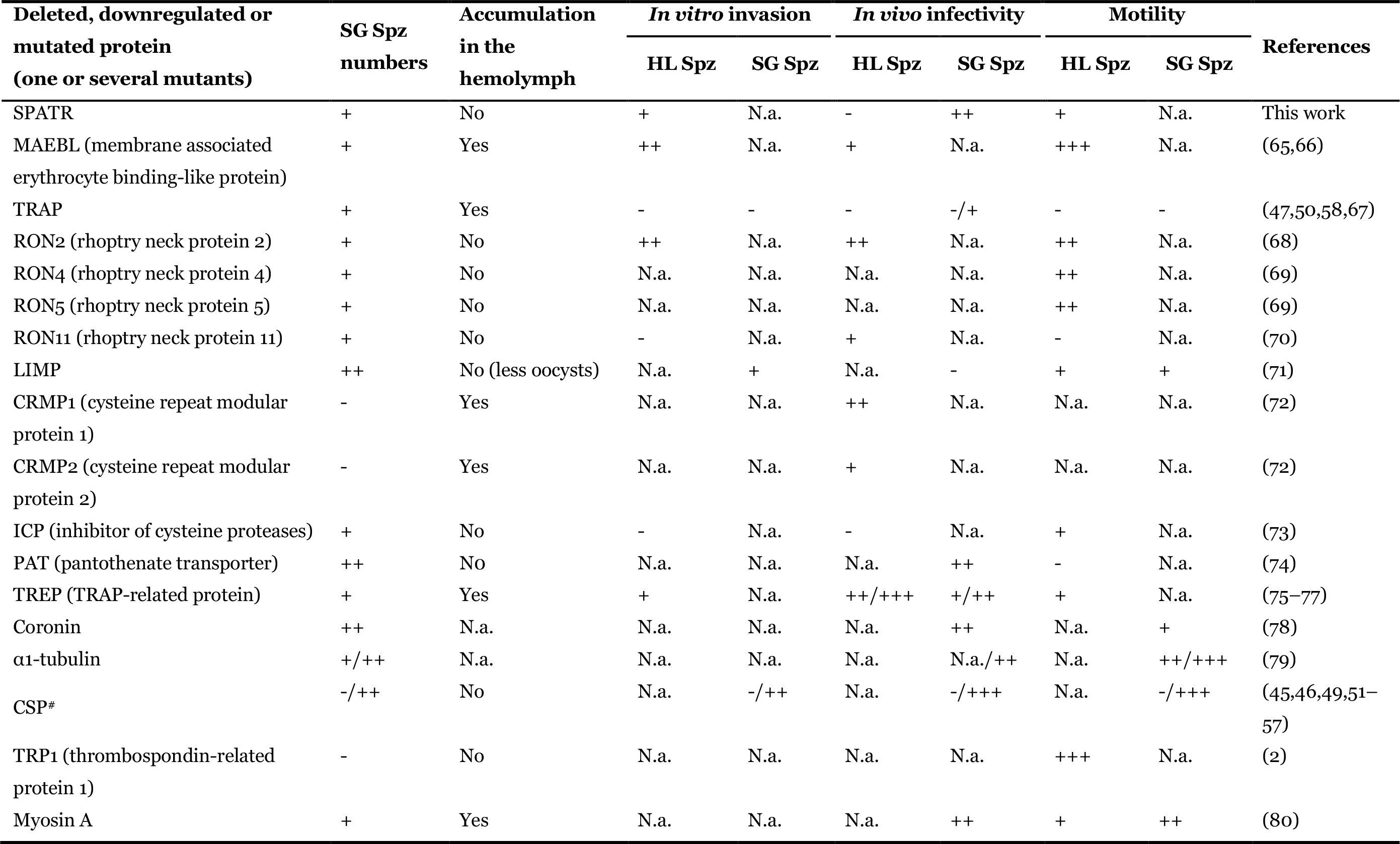
Phenotypic classification of mutant Plasmodium spp. lines displaying defective salivary gland invasion. -: abrogation of the process; +: severe defect; ++: mild defect; +++: no defect; ^#^: includes studies where antibodies and peptides were used to study salivary gland invasion; HL: hemolymph. N.a.: not applicable or addressed; SG: salivary gland; Spz: sporozoite.

